# Memory specific to temporal features of sound is formed by cue-selective enhancements in temporal coding enabled by inhibition of an epigenetic regulator

**DOI:** 10.1101/2021.03.31.437889

**Authors:** Elena K. Rotondo, Kasia M. Bieszczad

## Abstract

Recent investigation of memory-related functions in the auditory system have capitalized on the use of memory-modulating molecules to probe the relationship between memory and its substrates in auditory system coding. For example, epigenetic mechanisms, which regulate gene expression necessary for memory consolidation, are powerful modulators of learning-induced neuroplasticity and long-term memory formation (LTM). Inhibition of the epigenetic regulator histone deacetylase 3 (HDAC3) promotes LTM that is highly specific for *spectral* features of sound. The present work demonstrates for the first time that HDAC3 inhibition also enables memory for *temporal* features of sound. Rats trained in an amplitude modulation (AM) rate discrimination task and treated with a selective inhibitor of HDAC3 formed memory that was unusually specific to the AM rate paired with reward. Unusually sound-specific memory revealed behaviorally was associated with a signal-specific enhancement in temporal coding in the auditory system: stronger phase-locking that was specific to the rewarded AM rate was revealed in both the surface-recorded frequency following response (FFR) and auditory cortical multiunit activity in rats treated with the HDAC3-inhibitor. Furthermore, HDAC3inhibition increased trial-to-trial cortical response consistency (relative to naïve and trained vehicle-treated rats) that generalized across different AM rates. Stronger signal-specific phase-locking correlated with individual behavioral differences in memory specificity for the AM signal. Together, these findings support that epigenetic mechanisms regulate activity-dependent processes that enhance discriminability of sensory cues encoded into LTM in both spectral and temporal domains, which may be important for remembering spectrotemporal features of sounds, e.g., as in human voices and speech.

**SIGNIFICANCE STATEMENT:** Epigenetic mechanisms have recently been implicated in memory and information processing. Here, we use a pharmacological inhibitor of histone deacetylase 3 (HDAC3) in a sensory model of learning to reveal, for the first time, its ability to enable unusually precise memory for amplitude modulated sound cues. In so doing, we uncover neural substrates for memory’s “specificity” for temporal sound cues. Memory specificity was supported by auditory cortical changes in temporal coding, including greater response consistency and stronger phase-locking. HDAC3 appears to regulate effects across domains that determine specific cue saliency for behavior. Thus, epigenetic players may gate how sensory information is stored in long-term memory and can be leveraged to reveal the neural substrates of sensory details stored in memory.

## 3.1. INTRODUCTION

In humans and other species, precise representation of temporally modulated auditory signals is critical for identification of and discrimination between sounds used for communication. Individual differences in response timing, response consistency, and magnitude of phase-locked activity of the sound-evoked neural responses are associated with auditory behavioral ability (Hornickel et al 2009; Anderson et al., 2011; Anderson et al., 2012; Hornickel et al., 2012; Anderson et al. 2013; Centanni et al., 2014; Strait et al., 2014; Kraus et al., 2014; Tierney, Kirzman, & Kraus, 2015; Omote, Jasmin, & Tierney, 2017; Thompson et al., 2017; White-Schwoch et al., 2017; Otto-Meyer et al., 2018; von Trapp et al., 2016), and can be enhanced by learning (Beitel et al., 2003; Bao et al., 2004; Hornickel et al., 2012; Kraus et al., 2014; Tierney, Krizman, & Kraus, 2015; von Trapp et al., 2016). Indeed, experience can engender stimulus-specific neural enhancements for coding the temporal features of behaviorally relevant sounds (Bao et al., 2004; Song et al., 2008; Strait et al., 2012).

This study aimed to identify the auditory system substrates of newly formed memory that is specific to temporal features of sound. By leveraging a known memory-modulating molecule known to regulate the formation of unusually sound-specific memory by epigenetic control over consolidation processes—histone deacetylase 3 (HDAC3) (Stefanko et al., 2009; Malvaez et al., 2010, Gervain et al., 2013 Hitchcock et al., 2019; Bieszczad et al., 2015; Phan et al., 2017; Shang et al, 2019), its effects on temporal neural coding for temporal features of sound revealed novel relationships between individual differences in behavioral ability and their neural substrates in the auditory system. Rats were trained to discriminate among amplitude modulation (AM) rates either with or without treatment with a pharmacological HDAC3-inhibitor, which has been shown in *spectral* tasks to facilitate the formation of highly specific memory for acoustic frequency, with both cortical and subcortical sequelae (Bieszczad et al., 2015; Shang et al., 2019; Rotondo & Bieszczad, 2020; Rotondo & Bieszczad et al., 2021). AM was imposed on a broadband noise carrier, which virtually eliminated spectral cues available to rats and encouraged learning about AM rate. Following training, auditory cortical and subcortical responses were evoked by AM stimuli to determine learning-induced auditory system plasticity, its relationship to memory specificity for temporal cues, and whether either were facilitated by HDAC3 inhibition. This is the first study to address the role of HDAC3 in temporal information encoding for auditory memory and learning-induced auditory system plasticity. Furthermore, to complement intracortical recordings of auditory cortical responses, we used the surface-recorded frequency following response as a read-out of auditory-system wide integration changes in the representation of temporal sound cues as induced by learning and, for the first time, as modified by HDAC3 inhibition. Indeed, facilitating cortical and subcortical integration/coordination of experience-dependent enhancements for behaviorally significant features of sound might enable the encoding of persistent and vivid memories integral to hearing, speech, language, reading and musical abilities. Here, HDAC3-inhibition is applied to reveal how its effects extend to temporal forms of auditory processing that promote the precise storage of sensory details into associative memory that enable discriminative learned behavior.

## 2. METHODS

### 2.1 Subjects

A total of 21 adult male Sprague-Dawley rats (275-300 g on arrival; Charles River Laboratories, Wilmington MA) were used (RGFP966: n= 8; vehicle: n = 8; naïve: n = 5) in behavioral and electrophysiological procedures. In sum, these rats represent 3 separate groups: (1) treated-vehicle: rats received vehicle injections during amplitude modulation rate discrimination training, (2) treated-RGFP966: rats that received RGFP966 injections during amplitude modulation rate discrimination training, and (3) naïve: rats did not receive training in the auditory task and were used exclusively for baseline comparison of cortical electrophysiology. All animals were individually housed in a colony room with a 12-hour light/dark cycle. Throughout behavioral procedures, rats were water-restricted, with daily supplements provided to maintain at ∼85% free-drinking weight. All procedures were approved and conducted in accordance with guidelines by the Institutional Animal Care and Use Committee at Rutgers, The State University of New Jersey.

### 2.2 Behavioral Procedures and Analysis

All behavioral sessions were conducted in instrumental conditioning chambers within a sound-attenuated box. All subjects initially learned how to press a lever for water reward in five ∼45-minute bar-press shaping sessions before beginning the experimental timeline shown in Figure 1. This phase of training assured that all animals could acquire the procedural aspects of the task (i.e., bar-pressing for rewards) before any sounds were introduced.

**Figure 1.**
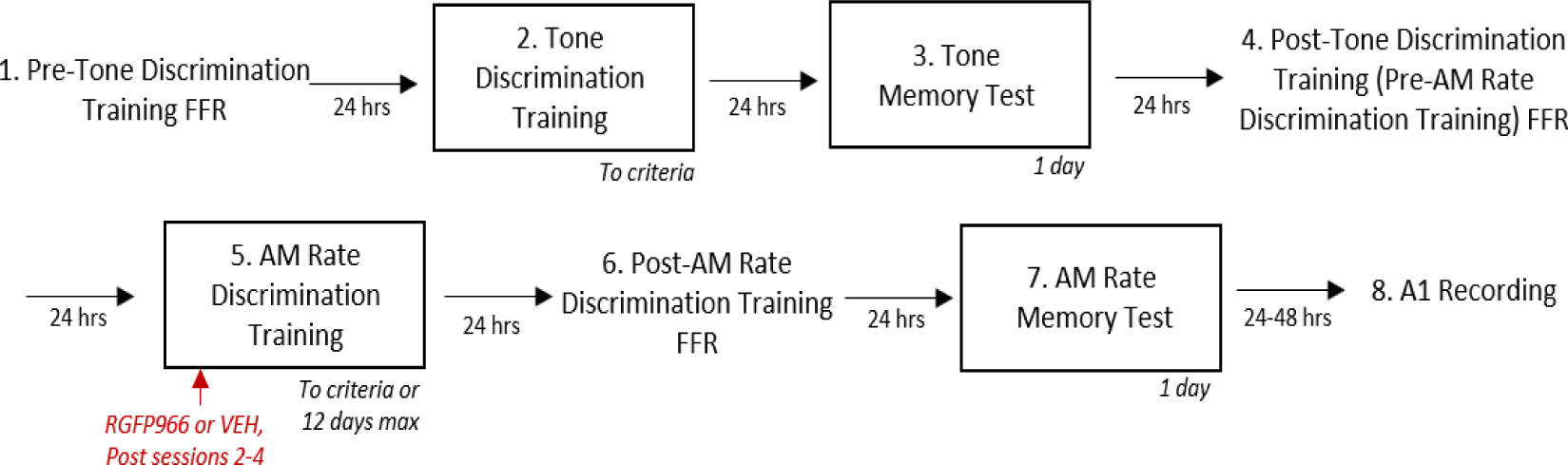
Experimental timeline. Rats underwent four behavioral phases (indicated in boxes): tone discrimination training, followed by a tone memory test, followed by AM rate discrimination training, followed by an AM rate memory test. Twenty-four hours prior to tone discrimination training, AM rate discrimination training, and the AM rate memory test, an AM noise-evoked frequency following response (FFR) was recorded. Twenty-four to forty-eight hours after the AM rate memory test, tone-and AM-noise evoked responses in the primary auditory cortex were recorded.

Next, rats received ***tone-tone discrimination training (TTD).*** The purpose of tone frequency-discrimination training was to establish that animals could perform the procedural aspects of the task under discrimination conditions (i.e., go vs. no-go). Further, it offers a way of measuring individual tendencies to form general or specific memories in a sound dimension other than the temporal cues under study and without any treatment with drugs.

Rats learned to respond to a 5 kHz pure tone S+ stimulus (8s, 70 dB) for operant reward and to withhold responding from a 9.8 kHz pure tone S-stimulus (8s, 70 dB). Responses to the S-or during the silent intertrial interval resulted in a visual error signal and triggered an 8 s time out that extended the time until the next sound trial. ***TTD*** training accomplished two goals. First, rats learned to bring operant behavior under control of sound cues (rather than silence). Second, rats learned the structure of a discrimination task. ***TTD*** training continued until rats reached criteria performance for two consecutive days (or a maximum of 20 training sessions). Criteria were (1) greater than 70% of all responses occurring in the presence of sound (vs. silence), (2) greater than or equal to 90% response rate to S+ trials, and (3) less than or equal to a 25% respond rate to S-trials.

Twenty-four hours following the following the final TTD training session, rats were given a ***Tone Memory Test*.** During the ***Tone Memory Test***, 8 pure tone frequencies were presented 12 times each in a pseudo-random order: 3.1, 4.0, 5.0, 6.2, 8.0, 9.8, 12.3, and 16.0 kHz. No responses were reinforced. The purpose of this test was to determine which animals were predisposed to signal-specific memory to increase experimental control prior to drug treatment manipulation.

To create performance matched groups for the remainder of the experiment, Measures of performance during ***TTD*** and of memory specificity as revealed by the ***Tone Memory Test*** were used to create performance matched groups for the remainder of the experiment (see *2.6.1 Behavioral Statistical Analysis)*.

Forty-eight hours following the ***Tone Memory Test***, rats began ***amplitude modulate (AM) rate discrimination training***. AM broadband noise (modulation depth: 100%; noise spectrum: 800-12,000 Hz; 8s, 70 dB) was presented 2 different modulation rates: 18.5 Hz (S+) and 155 Hz (S-). Each sound trial was separated by a variable ITI (mean = 15 s; range = 5-25 s). Rats learned to respond to the S+ for a reward. Bar presses to the S- or the silent ITI triggered an error signal and an 8 s time-out period. All rats were trained to performance criteria (or a maximum of 12 training sessions). Criteria were (1) greater than 70% of all responses occurring in the presence of sound (vs. silence), (2) greater than or equal to 90% response rate to S+ trials, and (3) less than or equal to a 25% respond rate to S-trials. ***AM rate discrimination training*** performance was calculated and statistically analyzed as described for ***tone discrimination training*** above.

Forty-eight hours following the final AM rate discrimination training session, rats were tested in an ***AM Rate Memory Test*** to reveal memory specificity for signal AM rates. During the memory test, 8 AM rates were presented 12 times each in a pseudo-random order: 4.1, 8.5, 18.5, 40, 79, 155, 307, and 625 Hz. No responses were reinforced.

### 2.3 HDAC3 Manipulation

The pharmacological HDAC3 inhibitor RGFP966 was used alter molecular mechanisms of auditory memory formation induced by learning (Bieszczad et al., 2015). Rats were assigned to either the RGFP966 (n= 8) or vehicle (n = 8) prior to AM rate discrimination training such that groups are matched with respect to tone-tone discrimination performance and predisposition to memory specificity as revealed by the Tone Memory Test (as described in *2.2 Behavioral Procedures and Analysis*). Rats received 3 consecutive days of post-session injections on AM rate discrimination training days 2-4 of RGFP966 (10 mg/kg, s.c.) or vehicle (equated for volume) (dose established [Malvaez et al., 2013], and confirmed in auditory system function [Bieszczad et al., 2015]). Post-training pharmacological treatment confines manipulation to the memory consolidation period, while avoiding potential performance effects based on perception, motivation, or within-session learning. For the remainder of training sessions, all rats received post-session injections of saline (equated for volume) to ensure that any effect of the injection itself remained consistent throughout training until reaching performance asymptote.

### 2.4 Frequency Follow Response Recordings and Analysis

The surface-recorded frequency following response (FFR), which measures neural activity phase-locked to sound, was recorded 3 times in anesthetized rats (ketamine-xylazine, K: 90 mg/kg, X:11 mg/kg, i.p.) to determine learning-induced changes in subcortical processing of sound: (1) 24 hours prior to tone-tone discrimination training, (2) 24 hours prior to AM rate discrimination training, and (3) 24 hours following the final AM rate discrimination training session. All recordings were made in a recording chamber completely separate from the training chamber and while the animal was anesthetized, which is a completely different state and context than that used in training. Stimulus presentation and neural response recordings were carried out using BioSig RZ software (TDT Inc.). Evoked potentials were recorded using a three-electrode configuration, with subdermal needle electrodes (1 kΩ) positioned at the midline along the head (recording), immediately below the left pinna (reference), and the midline on the back of the neck (ground). Sound stimuli were 70 dB SPL, AM broadband noise presented to left ear from a speaker positioned 8 cm away. Four AM rates (18.5, 40, 79, and 155 Hz) were presented in a blocked format (1500 stimuli per block, each block repeated two times). Stimulus duration varied according to AM rate to complete 5 full modulation cycles (range: 32.26 – 270.27 ms). The presentation rate ranged from 2.4/s to 10.1/s. Recordings were lowpass filtered online at 3 kHz and highpass filtered online at 10 Hz.

### 2.5 Auditory Cortical Recording Procedure and Analysis

To determine changes in auditory cortical response to AM noise, electrophysiological recordings were obtained from anesthetized subjects (total n = 21 rats) (sodium pentobarbital, 50mg/kg, i.p.) in an acute, terminal recording session 24-48 hours following the AM Memory Test. Recordings were obtained from trained animals (vehicle, RGFP966) and a group of experimentally naïve animals that did not receive behavioral training. All recordings were in the same recording chamber as what was used to obtain FFRs, which was completely separate from the training chamber while the animals were in a completely different state and context than that used in training. Therefore, any differences in responses measures from trained vs. naïve rats are interpreted to be robust and lasting changes to the auditory system’s response to sound that is not context-dependent nor dependent on short-term arousal or attention processes during active task engagement. All recordings were performed inside a double-walled, sound attenuated room using a linear array (2 x 3) of parylene-coated microelectrodes (1-2 MΩ, 250 µm apart) targeted to the middle cortical layers (III-IV, 400-600 µm orthogonal to the cortical surface) of the right primary auditory cortex (A1). Multiple penetrations were performed across the cortical surface, with an average of 27.04 (SE = 2.48) sites identified as within A1 per animal.

Acoustic stimuli were presented to the left ear from a speaker positioned ∼10 cm from the ear. Two sets of sounds were used. The first set of sounds were 50 ms pure tones (1-9 ms cosine-gated rise/fall time) presented in a pseudorandom order (0.5-54.0 kHz in half-octave steps; 70 dB SPL; 10 repetitions) with a variable inter-stimulus interval an average of 700 +/-100 ms. This set will enabled a rough determination of frequency tuning. The second set was AM broadband noise (modulation depth: 100%; noise spectrum: 800 -12,000 Hz) presented in pseudorandom order (4.1, 8.5, 18.5, 40, 79, 155, 307, and 625 Hz modulation rates; and unmodulated noise; 70 dB SPL; 20 repetitions) with a variable inter-stimulus interval an average of 1600 +/- 300 ms. The duration of each stimulus was the time it takes to complete 10 full cycles of modulation (range: 16.0 - 2439.02 ms). The unmodulated noise had a duration of 580.71 ms, which represents the average duration of all AM stimuli, and was used to confirm that the recording sites were sound responsive.

Neural activity was amplified x1000 and digitized for subsequent off-line spike detection and analysis using custom Matlab© scripts. Recordings were bandpass filtered (0.3-3.0 kHz). Multiunit discharges were characterized using previously reported temporal and amplitude criteria (Elias et al., 2015). Acceptable spikes were designated as waveforms with peaks separated by no more than 0.6 ms and with a threshold amplitude greater than 1.5 (for the positive peak) and less than 2.0 (for the negative peak) x RMS of 500 random traces from the same recording on the same microelectrode for each site. Responses greater than +/-1.0 SEM of the spontaneous spike rate were considered true sound-evoked responses.

### 2.6 Experimental Design and Statistical Analysis

Group sizes are consistent with prior reports showing brain-behavior relationships in rodent models of auditory memory, including with the use of an HDAC3 inhibitor (Bieszczad & Weinberger, 2010; Rotondo & Bieszczad, 2020; Rotondo & Bieszczad, 2021). Measures of the frequency following response are with-in subject, providing additional power to detect significant differences. Finally, correlative data between measures of neural plasticity and learned behavior further validate the current findings. All statistical analyses were performed using SPSS software (IBM, Armonk NY).

#### 2.6.1 Behavioral Statistical Analysis

Sixteen adult male Sprague Dawley rats (RGFP966 n= 8; vehicle n = 8) were used in the behavioral experiment. To create performance matched groups prior to AM rate discrimination training, measures of performance during ***TTD*** and of memory specificity as revealed by the ***Tone Memory Test*** were calculated and equated among groups. Metrics used to equate ***TTD*** performance included: **(1)** number of training days to criteria, **(2)** response rate to S+ trials during the first two vs. last two training sessions, **(3)** response rate to S-trials during the first two vs. last two training sessions, **(4)** the difference in response rate to S+ vs. S-trials during the first two vs. last two training sessions, and **(5)** sound control (% of responses occurring to sound vs. silence) during the first two vs. last two training sessions. Metric (1) was analyzed with an independent samples t-test, while metrics (2-5) were analyzed using a mixed-model ANOVA with the factors *drug treatment condition* and *session*.

Metrics used to equate tone memory specificity revealed by the ***Tone Memory Test*** included: **(1)** Percent of responses to the S+ frequency – percent of response to the S-frequency, where positive values indicate greater responding to the S+ than S-frequency. **(2)** For the S+ frequency, two contrast measures were calculated as follows: (a) Percent of responses to S+ frequency – (average percent of response to the nearby novel frequencies, 4.0 kHz and 6.2 kHz) and (b) Percent of responses to S+ frequency – (average percent of responses to distant novel frequencies, 3.1 and 8.0 kHz). Positive values indicate greater responding to the S+ than novel frequencies. **(3)** For the S-frequency, two contrast measures were calculated as follows: (a) Percent of responses to S-frequency – (average percent of response to the nearby novel frequencies, 8.0 and 12.3 kHz) and (b) Percent of responses to S-frequency– (average percent of responses to distant novel frequencies, 6.2 and 16.0 kHz). Negative values indicate less responding to the S-than novel frequencies. Additional metrics to examine group performance during the tone memory test included: **(1)** sound control (% of responses occurring to sound vs. silence) and **(2)** total number of bar presses to sound. All metrics were analyzed using an independent samples t-test between drug treatment groups.

To quantify memory specificity for the signal AM rates, contrast measures of relative responses to AM rate were determined in three ways. **(1)** Percent of responses to the S+ AM rate – percent of response to the S-AM rate, where positive values indicate greater responding to the S+ than S-AM rates. **(2)** For the S+ AM rate, two contrast measures were calculated as follows: (a) Percent of responses to signal AM rate – (average percent of response to the nearby AM rates, 8.5 Hz and 40 Hz) and (b) Percent of responses to signal AM rate – (average percent of responses to distant AM rates, 4.1 and 79 Hz). Positive values indicate greater responding to the S+ AM rate than novel AM rates. **(3)** For the S-AM rate, two contrast measures were calculated as follows: (a) Percent of responses to signal AM rate – (average percent of response to the nearby AM rates, 79 and 307 Hz) and (b) Percent of responses to signal AM rate – (average percent of responses to distant AM rates, 40 and 625 Hz). Negative values indicate less responding to the S-AM rate than novel AM rates. All metrics were analyzed using an independent samples t-test between drug treatment groups. In addition, a binomial test was used to test whether treated with an HDAC3 inhibitor shifted the proportion of animals that fell into the top vs. bottom half of memory specificity indices during the AM Rate Memory test compared to the Tone Memory Test.

#### 2.6.2 Frequency Following Response Recordings Statistical Analysis

FFRs were recorded at three time points for each of the 16 RGFP966-and vehicle-treated rats in the behavioral experiment (as specified in *2.4 Frequency Following Response Recording Procedures*). Valid responses evoked by 79 Hz and 40 Hz AM rates were unable to be obtained for 2 subjects (n = 1 vehicle-treated; n = 1 RGFP966-treated). Samples sizes used all FFR analyses are specified in Table 5.

Three analyses were used to determine changes in the FFR. In each case, values were compared pre-to-post training in each individual subject to measure learning-induced changes.

i. *Response Magnitude (dB)-* To determine response magnitude of phase-locked neural activity evoked by a given AM rate, a fast Fourier transform (FFT) was applied to the filtered averaged waveform (representing the average of both 1500-stimulus blocks, i.e. response to 3000 stimulus repetitions). Prior to performing the FFT, a Hanning window was applied to the signal to minimize the effects of DC offset on the resulting spectrum. Response magnitude (dB) of each response was calculated as follows:

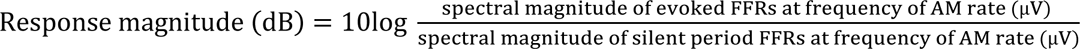

The difference in response magnitude as a function of learning was determined by subtracting pre-training response magnitude from post-training response magnitude for each individual.
ii. *Response Consistency (Fisher Z)*-Response consistency is a measure that captures variation in both timing and morphology for evoked responses. The average responses to each 1500-trial block of a given stimulus were used to derive measures of response consistency in the FFR. This method of analyzing response consistency has been previously validated, and published comparisons have shown a high correlation (r=0.98) among the first block vs. last block of trials and bootstrapping techniques based on single-trial FFR data (Hornickel & Kraus, 2013; Krizman & Kraus, 2019). The two sub-averages were correlated to produce an *r* value. Because the *r* value distribution is not normal, a fisher Z transformation was applied to the *r* values to increase the spread of the data. The difference in transformed r values at the pre-*vs.* post-training time point were calculated for each individual and for each AM rate.
iii. *Timing Jitter (ms)-* Timing jitter measures trial-to-trial variability in neural timing independently from morphology. The average responses to each 1500-trial block of a given stimulus were used to derive measures of jitter in the FFR. The two sub-averages were cross-correlated to determine the timing lag that produces the highest *r* value. The difference in the absolute value of the timing lag was calculated at the pre vs. post-training time pot for each individual and for each AM rate.

FFR data were analyzed using a series of Holm-Bonferroni corrected t-tests. Single-sample t-tests were used to determine whether a particular aspect of the FFR (response magnitude, consistency, or jitter) was significantly altered over the course of learning. Independent samples t tests were used to determine whether the change in a particular aspect of the FFR was different among the drug treatment groups.

One caveat in the present design is that the FFR will likely reflect activity of both cortical and subcortical sources for the slower S+ AM rate (18.5 Hz), whereas responses evoked by the faster AM rates will predominantly be of subcortical origin (Joris et al., 2004; Anderson et al., 2006; Fitzpatrick et al., 2009; Coffey et al., 2016). Therefore, all neural responses evoked by the S+ in this study will include a cortical component, precluding the ability to localize plasticity to only subcortical (vs. cortical) levels of the auditory system. Nonetheless, the design does provide the opportunity to validate our prior cortical findings using multiple recording methods and neural generators. On the other hand, the design does enable the observation of coordinated plasticity across cortical and subcortical levels for the faster S-AM rate (155 Hz).

#### 2.6.3 Auditory Cortical Recordings Statistical Analysis

Auditory cortical responses were compared among three groups: RGFP966-treated (n = 8), vehicle-treated (n=8), and behaviorally naïve (n = 8) rats. For group analyses, individual recording sites were treated as individual observations. Sample sizes for each analysis are reported in Tables 6-9. For each recording site, tone-evoked spike rate (spikes/s) were calculated by subtracting spontaneous spiking (40 ms window prior to tone onset) from evoked-spiking within a 40 ms response-onset window (6-46 ms after each tone onset). Tone-evoked neural activity was used to determine tonotopy to identify recording cites within the primary auditory cortex (A1). Best frequency (BF; the frequency to which the site responds to best at a given sound level-i.e. 70 dB) of each recording site was determined using evoked spike rate as a function of frequency. Sites that exhibit the characteristic progression from low-high BF along the posterior-anterior axis were classified as A1 and used for subsequent analyses. A chi-square test was used to test for group differences in the distribution of recording site BFs between groups.

Among recording sites within A1, AM noise-evoked multi-unit activity (Fig. 2) was used to learning-induced changes in AM encoding by analyzing several measures:

i. *Phase Locking:* Vector strength (VS) and the Rayleigh statistic (RS) were used to determine how well evoked activity was time-locked to the AM rates (Bao et al., 2004; Yao & Sanes, 2018; Yao & Sanes, 2021) For each AM rate, evoked neural responses from a given recording site were analyzed from 12 ms post-sound onset (to account for neural transmission delays) to 12 ms post-sound offset across 20 stimulus repetitions. Durations were set to the time needed to complete 10 AM cycles. Note that each sound was of a different duration, to account for the different AM rates. VS was calculated as across the evoked window as in previous studies (Goldberg and Brown, 1969; Bao et al., 2004; Yao & Sanes, 2018):

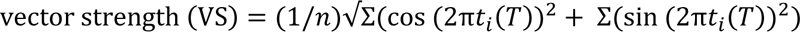

Here, t*i* (*i* = 1,2…n) is the time between the onset of the stimulus and the i*th* spike and *T* is the period the amplitude modulation. One-way ANOVAs, followed by Holm-Bonferroni corrected t-tests, were used to determine group differences in vector strength in evoked responses. The Rayleigh statistic, which estimates the significance of phase-locking while controlling for the total number of spikes, was calculated as in previous studies (Mardia, 1972; Bao et al., 2004; Yao & Sanes, 2018):

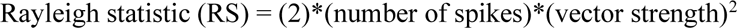

The critical values for the Rayleigh statistic are 5.991 corresponding to p<0.05 and 13.816 corresponding to p<0.001. The threshold value of 5.991 was used to categorize sites into phase-locking or non-phase-locking responses to determine the proportion of recording sites that exhibited significant phase-locking by subjects and by treatment group. A binomial test was used to test for significant differences in the proportion of phase-locked sites recorded from vehicle-treated or drug-treated rats, each independently compared to a group of untrained naïve rats. In a subset of analyses of vector strength, only responses from phase-locked sites, as determined by the Rayleigh statistic, were used. One rat did not have any sites with significant phase-locking, and was thus excluded from these analyses.
ii. *Response Consistency:* To determine within-stimulus response consistency in the AM evoked neural responses, we used a *k-means* clustering approach over 20 individual trials of a given AM rate for each recording site. This approach generated a multi-dimensional hyperplane that represented spike counts across 5 ms time bins in the same evoked response period as used previously (12 ms after sound onset to 12 ms after sound offset) for each individual stimulus trial over 20 repetitions. The average spike count in time bins across trials was used to generate the centroid in the *k-means* cluster. The sum of distances from each repetition in the cluster to the centroid (“SUMD” in the *kmeans* Matlab© function) was used to represent the consistency of the AM evoked cortical response to any unique stimulus. *Lower* SUMD values indicate *greater* response consistency (i.e., and lower response variability). One-way ANOVAs, followed by Holm-Bonferroni-corrected t-tests, were used to determine group differences. Corrected p-values are reported. Further, Pearson correlations were used to test for correlative relationships between cortical response consistency and phase-locking.

**Figure 2.**
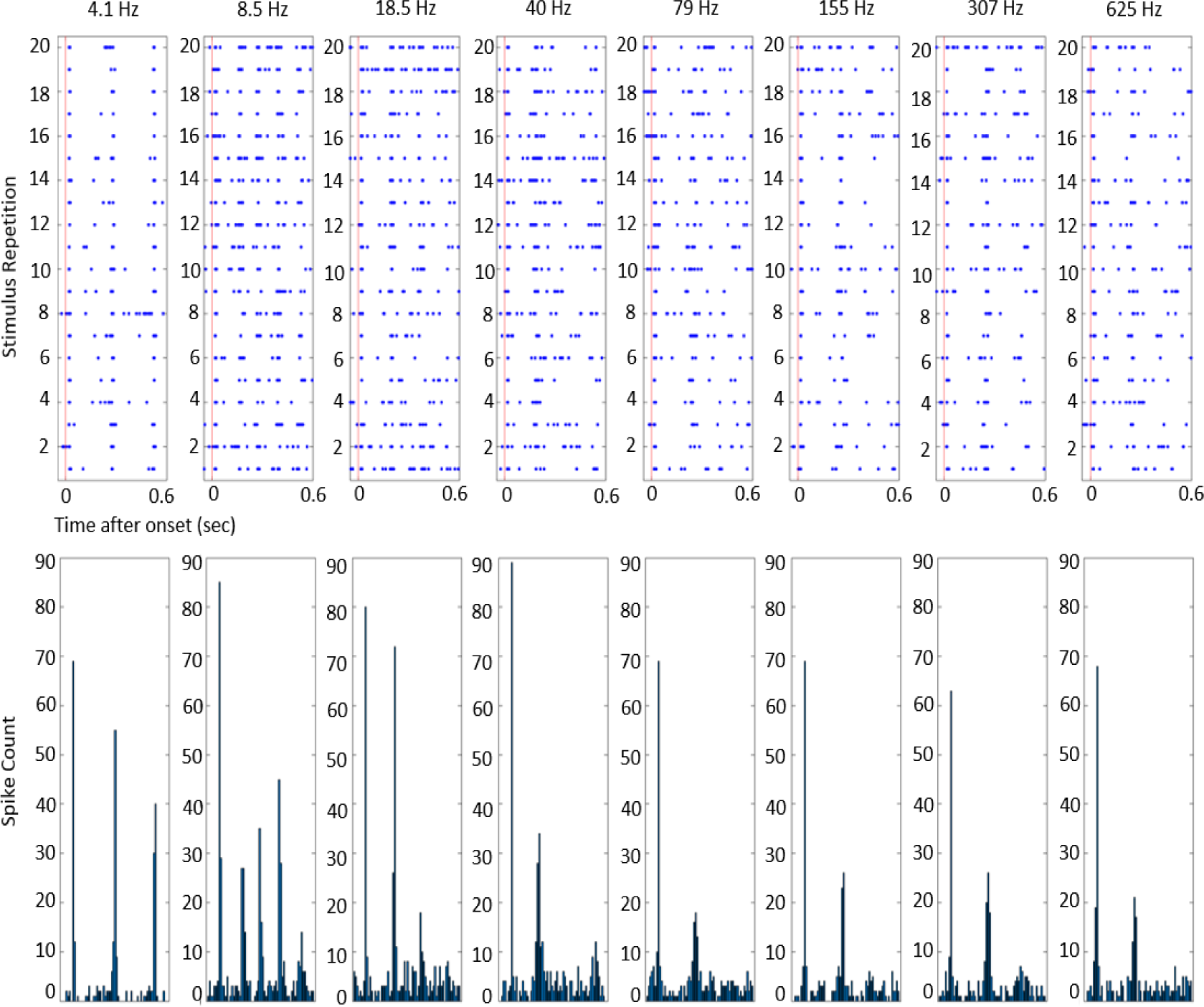
AM noise-evoked auditory cortical responses. Eight different AM rates, ranging from 4.1 to 625 Hz, were used to evoked auditory cortical responses. Each AM rate was presented 20 times in a pseudorandom order, with responses to each trial represented in the spike rasters in the upper panels. The lower panels display the peri-stimulus time histogram (from 0 to 0.6 seconds post-sound onset) collapsed across all trials for each AM rate. Note the periodic activity evoked by slower AM rates that tapers off with faster AM rates.

#### 2.6.4 Brain-Behavior Correlative Data

To limit the number of statistical comparisons, we set the *a priori* criterion that brain-behavior Pearson correlations would be pursued only for brain measures that showed signal-specific, learning-induced plasticity. The number of subjects available for each correlation (determined by subject attrition in neural measures, as described in sections *2.6.2 Frequency Following Response Recordings Statistical Analysis and 2.6.3 Auditory Cortical Recordings Statistical Analysis*) is detailed in Table 10.

Measures used for Pearson correlations are as follows. For behavioral measures, the percent of responses to the 18.5 Hz S+ was used as an index of memory specificity, as it was highly correlated with other indices of memory specificity (% BPs to S+ vs. Δ % BPs to S+ vs. S-: r = 0.981, p <0.00001; vs. Δ % BPs to S+ vs. nearby neighbors: r = 0.987, p <0.00001; vs. Δ % BPs to S+ vs. distant neighbors: r = 0.976, p < 0.00001). For FFR data, the change in response magnitude over the course of AM rate discrimination training in FFRs evoked by the 18.5 was used for each subject. For auditory cortical data, an average value was computed for each subject for vector strength, the Rayleigh statistic, and percent of phase-locked sites.

## 3 RESULTS

### 3.1 Groups perform identically during tone discrimination training and memory testing, prior to administration of the HDAC3 inhibitor

All rats were first trained in a tone discrimination task to bring operant responding under the control of sound and teach rats the procedural aspects of a discrimination task. Subsequently, rats underwent a Tone Memory Test to determine the specificity with which they remembered the rewarded S+ (5.0 kHz) and unrewarded S-(9.8 kHz) tones. Behavioral performance during both training and testing was used to create behaviorally equivalent groups prior to administration of the HDAC3 inhibitor in the following behavioral training phase (Tables 1,2; Fig. 3).

**Figure 3.**
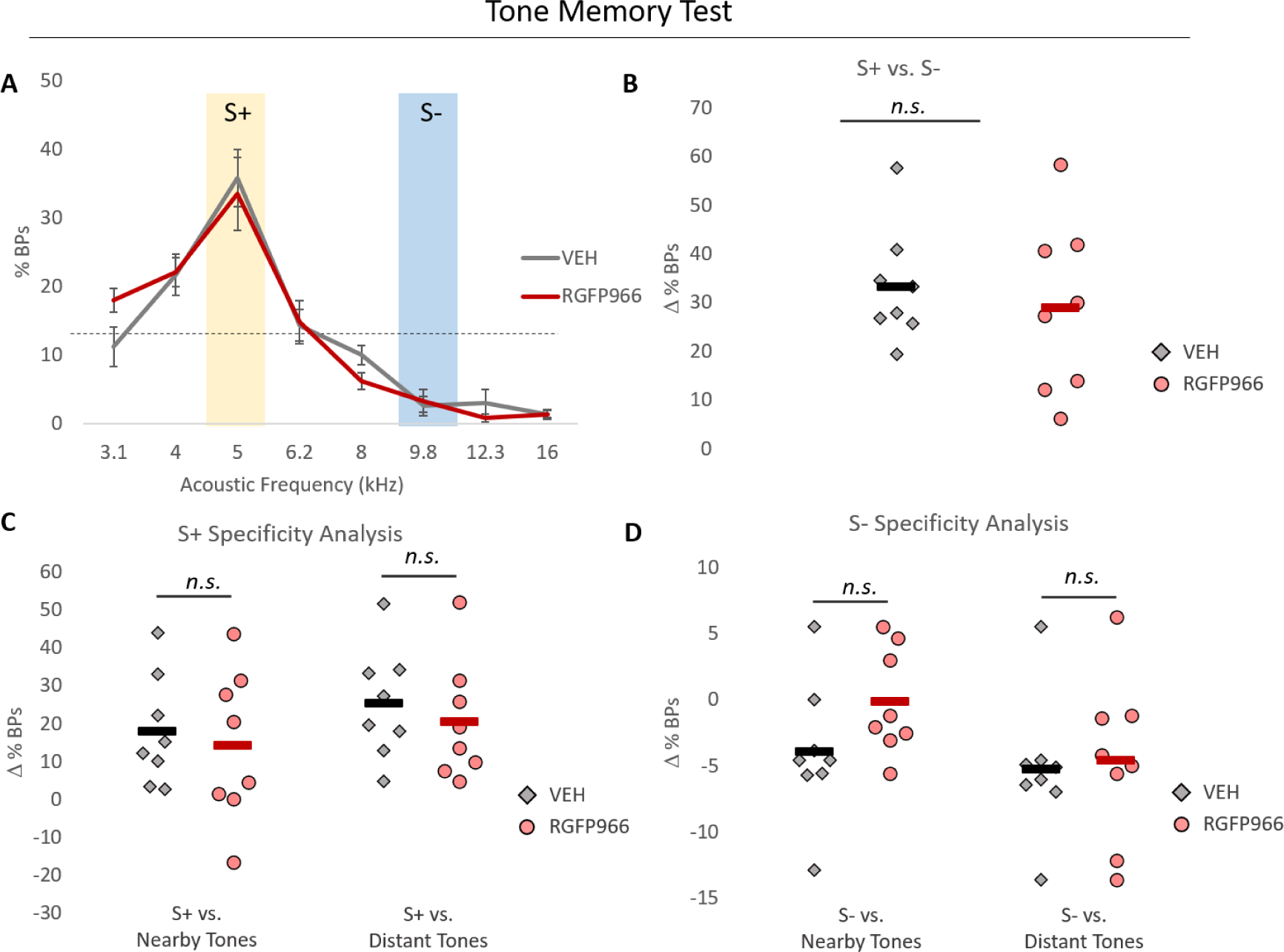
Prior to drug treatment, groups are matched for frequency-specificity of memory in the Tone Memory Test. (A) Groups exhibit similar response distributions across frequencies during the Tone Memory Test. The dashed line represents the Memory Test gradient if responses were equally distributed among the frequencies, which would indicate a completely generalized memory. The shape of the response distribution was quantified using relative measures of responding to S+, S-, and novel tone frequencies. These measures revealed no group differences in discrimination of the S+ frequency relative to (B) the S- or (C) the nearby and distant novel tone neighbors. (D) There were no group differences in discrimination of the S-tone frequency from its nearby and distant novel tone neighbors. Dots represent individual subjects. Bars represents the group mean.

**Table 1.** Tone discrimination training performance metrics for subsequent drug treatment groups. This table displays performance metrics for vehicle- and RGFP966-treated rats during the first two vs. last two training sessions, including sound control (% of total responses that occurred to sound, as opposed to silence), S+ and S-response rate (percent of S+ or S-trials with at least one response), and the difference between S+ and S- response rate. Data is displayed as M ± SE.

| | | Sound Control | S+ Response Rate (%) | S- Response Rate (%) | $\Delta$ S+ - S- Response Rate (%) |
| --- | --- | --- | --- | --- | --- |
| First Two Sessions | VEH (n=8) | 15.25 ± 1.72 | 26.03 ± 6.21 | 28.78 ± 7.18 | -2.68 ± 1.98 |
| | RGFP966 (n=8) | 14.25 ± 1.69 | 24.64 ± 3.65 | 26.02 ± 3.21 | -2.13 ± 2.83 |
| Last Two Sessions | VEH (n=8) | 90.77 ± 1.09 | 98.56 ± 0.33 | 18.01 ± 1.40 | 80.47 ± 1.29 |
| | RGFP966 (n=8) | 88.99 ± 1.80 | 97.88 ± 0.53 | 15.16 ± 1.34 | 83.84 ± 1.81 |

**Table 2.** Performance during the Tone Memory Test. This table displays relative measures of responding to tone frequencies presented during the Tone Memory Test for subsequent vehicle-and RGFP966-treated rats. On average, both groups responded *more* to the S+ than the S- and the neighboring novel tones. Both groups also responded *less* to the S-than the neighboring novel tones. Data are displayed as M ± SE.

| | $\Delta$ % BPs to S+ vs. S- | $\Delta$ % BPs to S+ vs. Nearby Tones | $\Delta$ % BPs to S+ vs. Distant Tones | $\Delta$ % BPs to S- vs. Nearby Tones | $\Delta$ % BPs to S- vs. Distant Tones |
| --- | --- | --- | --- | --- | --- |
| VEH (n=8) | 33.24 ± 4.17 | 17.79 ± 5.13 | 25.17 ± 5.16 | -3.94 ± 1.85 | -5.26 ± 1.85 |
| RGFP966 (n=8) | 28.80 ± 6.26 | 14.10 ± 7.07 | 20.50 ± 5.55 | -0.13 ± 1.42 | -0.22 ± 0.82 |

Based on the subsequently assigned drug treatment conditions, the number of training days to reach criteria was equivalent among rats in the RGFP966 treatment group (M = 12.87, SE = 1.31) and the vehicle treatment group (M = 11.75, SE = 1.26) (t(14) = -0.616, p = 0.547). Overall, rats showed significant improvement on each performance metric between the first two and last two training sessions (sound control: F(1,14) = 3030.713, p < 0.001; S+ response rate: F(1,14) = 379.286, p<0.001; S-response rate: F(1,14) = 6.688, p = 0.022; differences in S+ vs. S-response rate: F(1,14) = 1420.594, p < 0.001). However, no metric showed a significant main effect for drug condition (sound control: F(1,14) = 0.590, p = 0.455 ; S+ response rate: F(1,14) = 0.088, p = 0.771; S-response rate: F(1,14) = 0.608, p = 0.448; difference in S+ vs. S-response rate: F(1,14) = 1.117, p = 0.308), and no metric showed a *significant training session x drug condition* interaction (sound control: F(1,14) = 0.081, p = 0.780; S+ response rate: F(1,14) = 0.009, p = 0.925; S-response rate: F(1,14) < 0.000, p = 0.987; difference in S+ vs. S-response rate: F(1,14) = 0.396, p = 0.539). Group means and SEs are displayed in Table 1

Based on the subsequently assigned drug treatment conditions, independent samples t-tests revealed that memory specificity was identical among groups (Fig. 3; Table 2). Further, ranking each rat from greatest to smallest percentage of responses to the S+ revealed that 4 of the 8 rats (i.e., 50%) were in the top half were from each drug condition. Similarly, ranking each rat from greatest to smallest difference in the percentage of responses to the S+ vs. S-revealed that 4 of the 8 rats (50%) were from each drug condition. Additionally, other performance metrics were equivalent between groups, including sound control (RGFP966: M = 83.48, SE = 3.14; VEH: M = 88.93, SE = 1.84; t(14) = 1.465, p = 0.164) and the number of bar presses to sound (RGFP966: M = 37.25, SE = 5.55; VEH: M = 37.12, SE = 3.79; t(14) = -0.018, p = 0.985). Together, these behavioral results show that there was no *a priori* difference between animals later assigned to vehicle- or RGFP966-treatment groups in relevant measures of sound-evoked bar-press responding or in tendency to remember sound specifically (to the S+ sound frequency) or generally (across a *spectral* feature of sound: acoustic frequency). Therefore, subsequent differences in performance during AM rate discrimination training or the AM rate memory test can be attributed to the effects of HDAC3 inhibition, rather than procedural learning aspects of the task.

### 3.2 HDAC3 inhibition promotes memory specificity for AM rate

During AM rate discrimination training, rats received injections of either the HDAC3 inhibitor RGFP966 or vehicle immediately following sessions 2-4. RGFP966- and vehicle-treated rats did not differ in the number of number of days of AM rate discrimination training prior to the AM Memory Test (RGFP966: M = 11.62, SE = 0.37; VEH: M = 11.62, SE = 0.37; t(14) = 0, p > 0.999). Overall, rats showed improved performance on all metrics between the first two and final two training sessions (sound control: F(1,14) = 24.263, p < 0.001; S+ response rate: F(1,14) = 15.056, p = 0.002; S-response rate: F(1,14) = 13.860, p = 0.002; difference in S+ vs. S-response rate: F(1,14) = 77.937, p < 0.001). However, there was no main effect of drug treatment condition on any metric (sound control: F(1,14) = 0.001, p = 0.979; S+ response rate: F(1,14) = 2.129, p = 0.167; S-response rate: F(1,14) = 0.236, p = 0.634; difference in S+ vs. S-response rate: F(1,14) = 0.701, p = 0.417 ), nor a significant *drug treatment condition x session* interaction (sound control: F(1,14) = 0.032, p = 0.860; S+ response rate: F(1,14) = 0.141, p = 0.713; S-response rate: F(1,14) < 0.001, p = 0.997; difference in S+ vs. S-response rate: F(1,14) = 0.099, p = 0.757) (See Fig 4, Table 3). Importantly, because groups did not differ in performance during AM rate discrimination training means that any subsequent behavioral differences during the AM rate memory test can be attributed to the remembered discriminability of AM stimuli alone.

**Figure 4.**
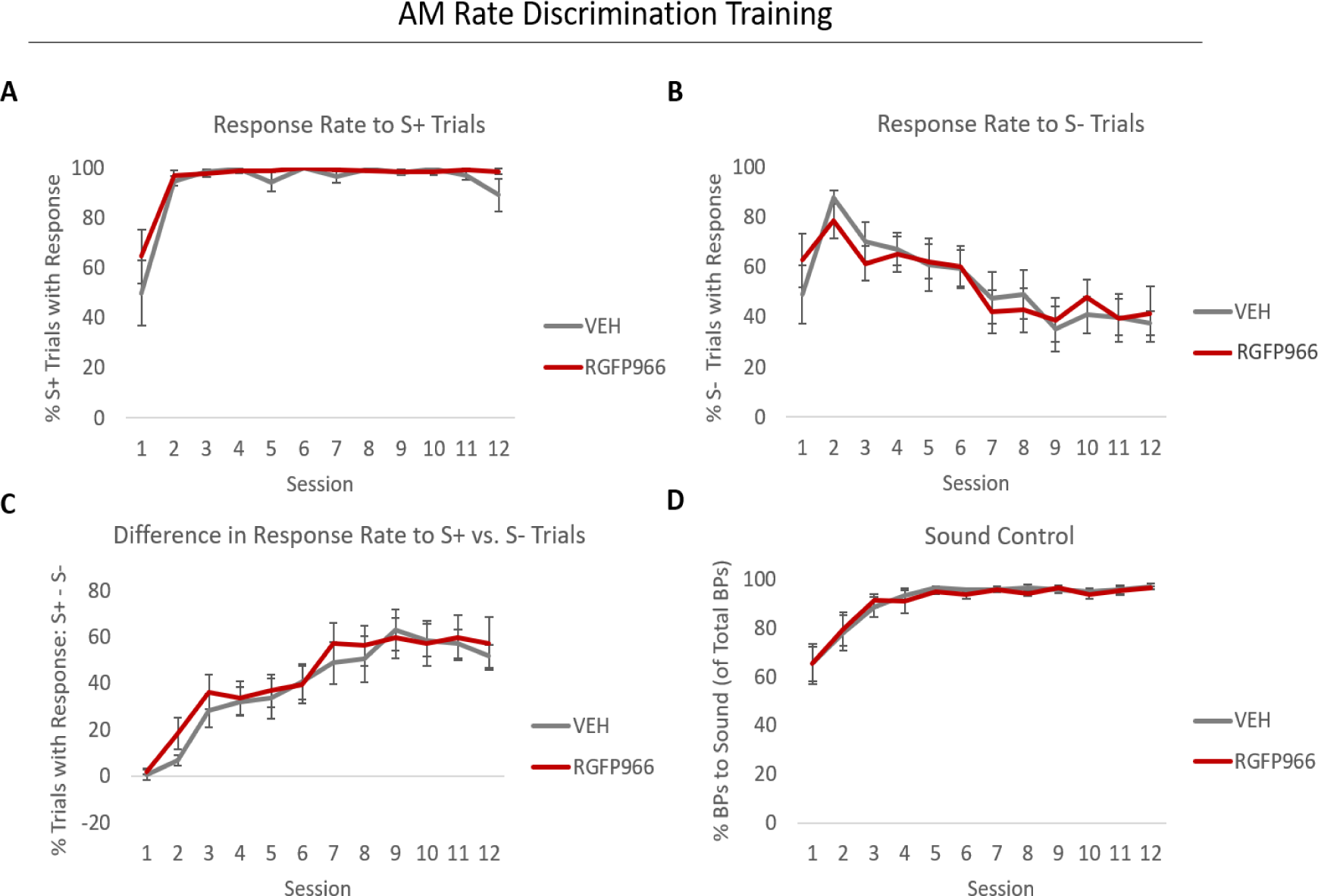
HDAC3 inhibition does not alter performance in the amplitude modulate rate discrimination task. Performance metrics are equivalent between groups treated with RGFP966 and vehicle, including (A) response rate to S+ trials, (B) response rate to S-trials, (C) difference in response rate to S+ vs. S-trials, and (D) sound control (the percent of response occurring to either sound vs. silence).

**Table 3.** AM rate discrimination training performance metrics for drug treatment groups. This table displays performance metrics for vehicle- and RGFP966-treated rats during the first two vs. last two training sessions, including sound control (% of total responses that occurred to sound, as opposed to silence), S+ and S-response rate (percent of S+ or S-trials with at least one response), and the difference between S+ and S- response rate. Data is displayed as M ± SE.

| | | Sound Control | S+ Response Rate (%) | S- Response Rate (%) | $\Delta$ S+ vs. S- Response Rate (%) |
| --- | --- | --- | --- | --- | --- |
| First Two Sessions | VEH (n=8) | 71.73 ± 6.84 | 72.29 ± 6.97 | 68.45 ± 5.82 | 3.84 ± 2.25 |
| | RGFP966 (n=8) | 61.27 ± 10.57 | 80.79 ± 5.43 | 70.66 ± 7.07 | 10.12 ± 3.43 |
| Last Two Sessions | VEH (n=8) | 96.41 ± 1.19 | 93.74 ± 3.57 | 36.04 ± 5.31 | 57.69 ± 5.53 |
| | RGFP966 (n=8) | 91.04 ± 4.59 | 98.44 ± 0.86 | 38.18 ± 8.93 | 60.26 ± 8.90 |

The AM rate memory test revealed that RGFP966-treated rats formed memory for the rewarded S+ AM rate with greater specificity than vehicle-treated rats (Fig. 5; Table 4). Moreover, ranking each rat based on the percentage of responses to the S+ revealed that 7 of 8 rats in the top half were treated with RGF966 (vs. 4 of 8 during the tone memory test; binomial test: p = 0.038). Similarly, 7 of 8 rats with the greatest difference in responding to the S+ vs. S- were treated with RGFP966 (vs. 4 of 8 during the tone memory test; binomial test: p = 0.038). Therefore, treatment with RGFP966 shifts the behavioral distribution toward greater memory specificity for the S+. In contrast, groups did not differ with respect to memory specificity for the unrewarded S- AM rate; responses to the S- stimulus during the memory test were equivalent (Table 4). Further, drug treatment groups did not differ on other performance metrics, including sound control measured by the percent of responses to either AM sound vs. responses during silent inter-trial intervals (RGFP966: M = 86.45, SE = 2.04; VEH: M = 82.74, SE = 4.32; t(14) = -0.774, p = 0.451) or number of bar presses to AM sounds (RGFP966: M = 39.37, SE = 10.91; VEH: M = 40.5, SE = 6.28; t(14) = 0.089, p = 0.930). Therefore, the main effect of RGFP966 on the AM task was to increase memory specificity for the AM rate paired with reward (S+), adding to existing evidence that effects of HDAC3i on memory specificity can occur independently from effects on rate of learning (Rotondo & Bieszczad, 2020; Rotondo & Bieszczad, 2021).

**Figure 5.**
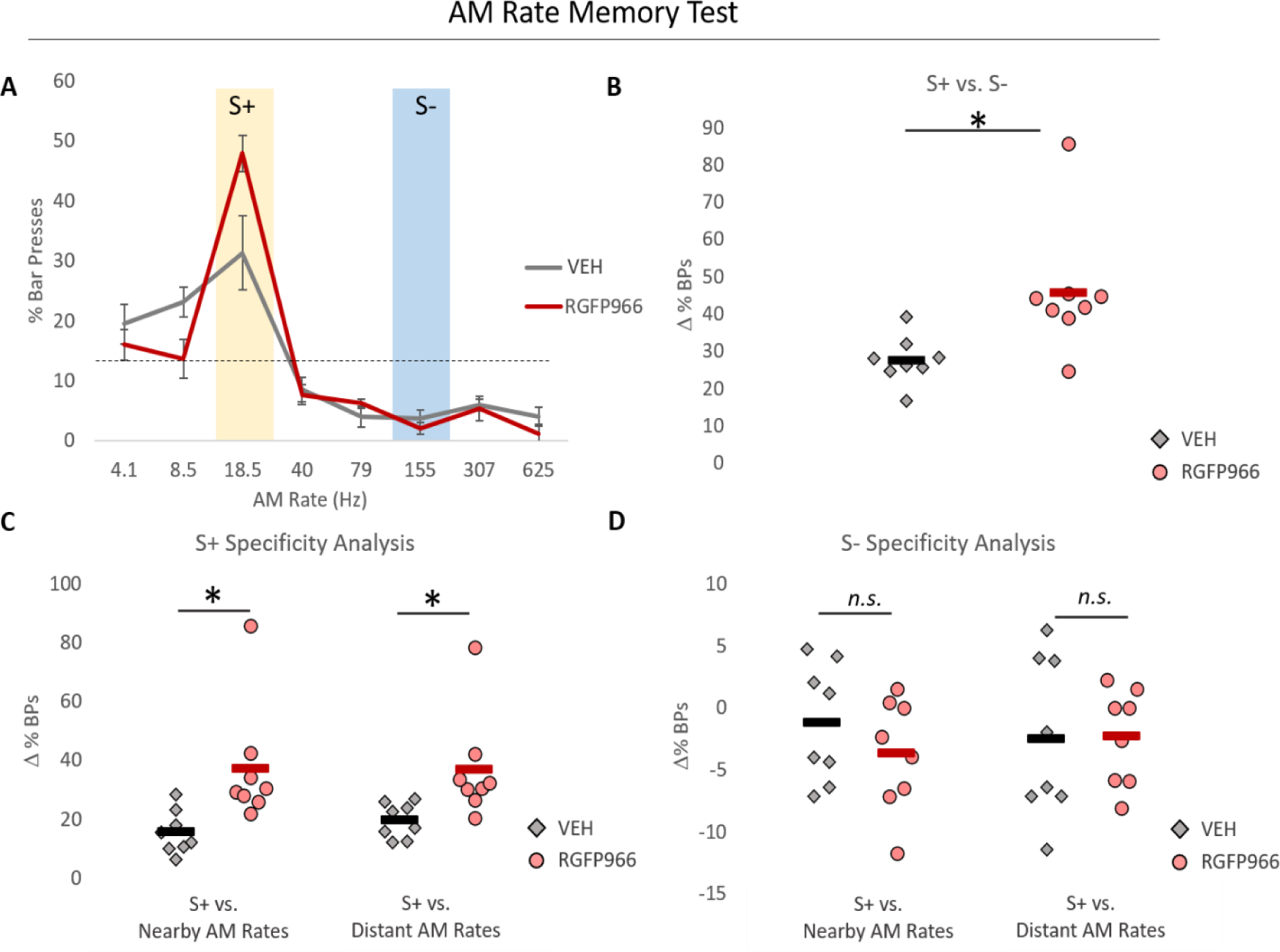
HDAC3 inhibition promotes memory specificity for AM rate. (A) RGFP966-treated rats exhibit an AM- rate specific response distribution with a sharper peak at the S+ AM rate compared with vehicle-treated rats. The dashed line represents the Memory Test gradient if responses were equally distributed among all AM rates, which would indicate a completely generalized memory. The shape of the response distribution was quantified using relative measures of responding to S+, S-, and novel AM rates. Compared with vehicle-treated rats, HDAC3 inhibited rats showed greater discrimination for the S+ AM rate from (B) the S- and (C) the nearby and distant novel AM rate neighbors. (D) There were no group differences in discrimination of the S- AM rate from its nearby and distant novel AM rate neighbors. Dots represent individual subjects. Bars represents the group mean.

**Table 4.** Performance during the AM Rate Memory Test. This table displays relative measures of responding to AM rates presented during the AM rate Memory Test for vehicle- and RGFP966-treated rats. On average, both groups responded *more* to the S+ than the S- and the neighboring novel tones. However, RGFP966-treated rats showed a greater response bias toward the S+ than vehicle treated rats. Both groups also responded *less* to the S- than the neighboring novel tones, with no drug treatment group differences. Data are displayed as M ± SE.

| | $\Delta$ % BPs to S+<br>vs. S- | $\Delta$ % BPs to S+<br>vs. Nearby AM<br>Rates | $\Delta$ % BPs to S+<br>vs. Distant AM<br>Rates | $\Delta$ % BPs to S-<br>vs. Nearby AM<br>Rates | $\Delta$ % BPs to S-<br>vs. Distant AM<br>Rates |
| --- | --- | --- | --- | --- | --- |
| VEH (n=8) | 27.59 ± 2.27 | 15.48 ± 2.60 | 19.58 ± 2.10 | -1.23 ± 1.69 | -2.51 ± 2.29 |
| RGFP966 (n=8) | **45.82 ± 6.17<sup>\*1</sup>** | **37.23 ± 7.25<sup>\*2</sup>** | **36.81 ± 6.34<sup>\*3</sup>** | -3.70 ± 1.60 | -2.32 ± 1.36 |

<sup>1</sup> t(14) = -2.277, p = 0.014
<sup>2</sup> t(14) = -2.819, p = 0.013
<sup>3</sup> t(14) = -2.576, p = 0.021

**Table 5.**
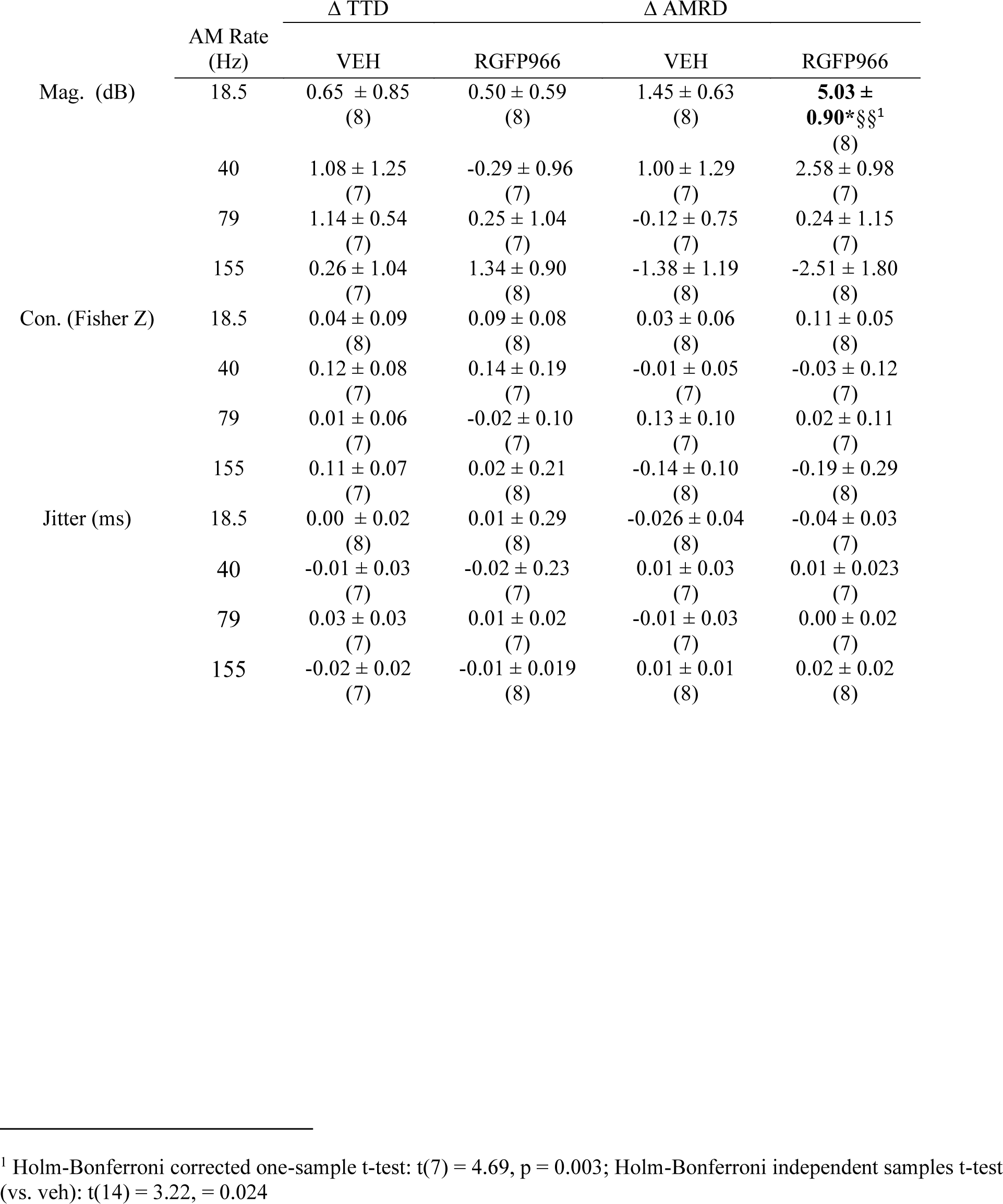
Changes in AM noise-evoked FFRs in vehicle- and HDAC3-treated animals. This table displays the change in FFR response magnitude (mag.), response consistency (con.), and timing jitter over the course of tone-tone discrimination training and over the course of AM rate discrimination training. All data are displayed as M ± SE, with sample sizes in parentheses. Significant differences are bolded. *indicates a difference from 0 (i.e. no change), *p<0.05. § indicates a difference vs. vehicle-treated animals, §§p<0.01

### 3.3 HDAC3 inhibition enhances response magnitude of FFRs evoked by the rewarded AM rate

The frequency-following response (FFR) is a non-invasive recording of neural activity that is phase-locked to periodic features of sound, including amplitude modulation. Neural generators of the FFR may vary with characteristics of the sounds used to evoke it (Coffey et al., 2016). Under present conditions, we assume that FFRs evoked by the 18.5 Hz AM sound have both cortical and subcortical components, while FFRs evoked by the remaining sound set (40, 79, and 155 Hz) are primarily of subcortical origin. Previous work has shown that the FFR may change with experience in a way that is selective for behaviorally relevant sounds (Song et al., 2008; Strait et al., 2012). The present study recorded AM noise-evoked FFRs (1) prior to tone-tone discrimination training, (2) post tone-tone discrimination training, and (3) post-AM rate discrimination training. Over the tone-tone discrimination training phase interval (i.e. prior to any experience with AM sounds), we predicted that there would be no change in AM noise-evoked FFRs. Over the AM rate discrimination training phase interval, it was predicted that memory specific to AM rate would be associated with changes in the FFR that are selective to the behaviorally relevant AM rates.

In support of the first prediction, characteristics of AM-evoked FFRs, including response magnitude, response consistency, and timing jitter, were stable over the course of tone-tone discrimination training (Table 5, Fig 6c). In addition, there were no differences in the change in these response characteristics between the subsequent drug treatment groups (see Table 5). Consistent with second prediction, over the course of AM rate discrimination training, there was a significant increase in response magnitude in FFRs evoked by the rewarded 18.5 Hz AM rate, but only in the RGFP966-treated group (Table 5, Fig. 6d). There were no changes in response magnitude in FFRs evoked by any other AM rate, including the unrewarded 155 Hz AM rate. Response consistency and timing jitter were stable over the course of AM rate discrimination training (Table 5). Therefore, the FFR reveals cue-selective enhancements in auditory processing due to learning. These effects are most prominent under conditions of unusually specific memory enabled by HDAC3 inhibition. Given the presumed neural generators of the 18.5 Hz-evoked FFR, results point to a potential cortical change in learning-induced processing of behaviorally relevant temporal cues.

**Figure 6.**
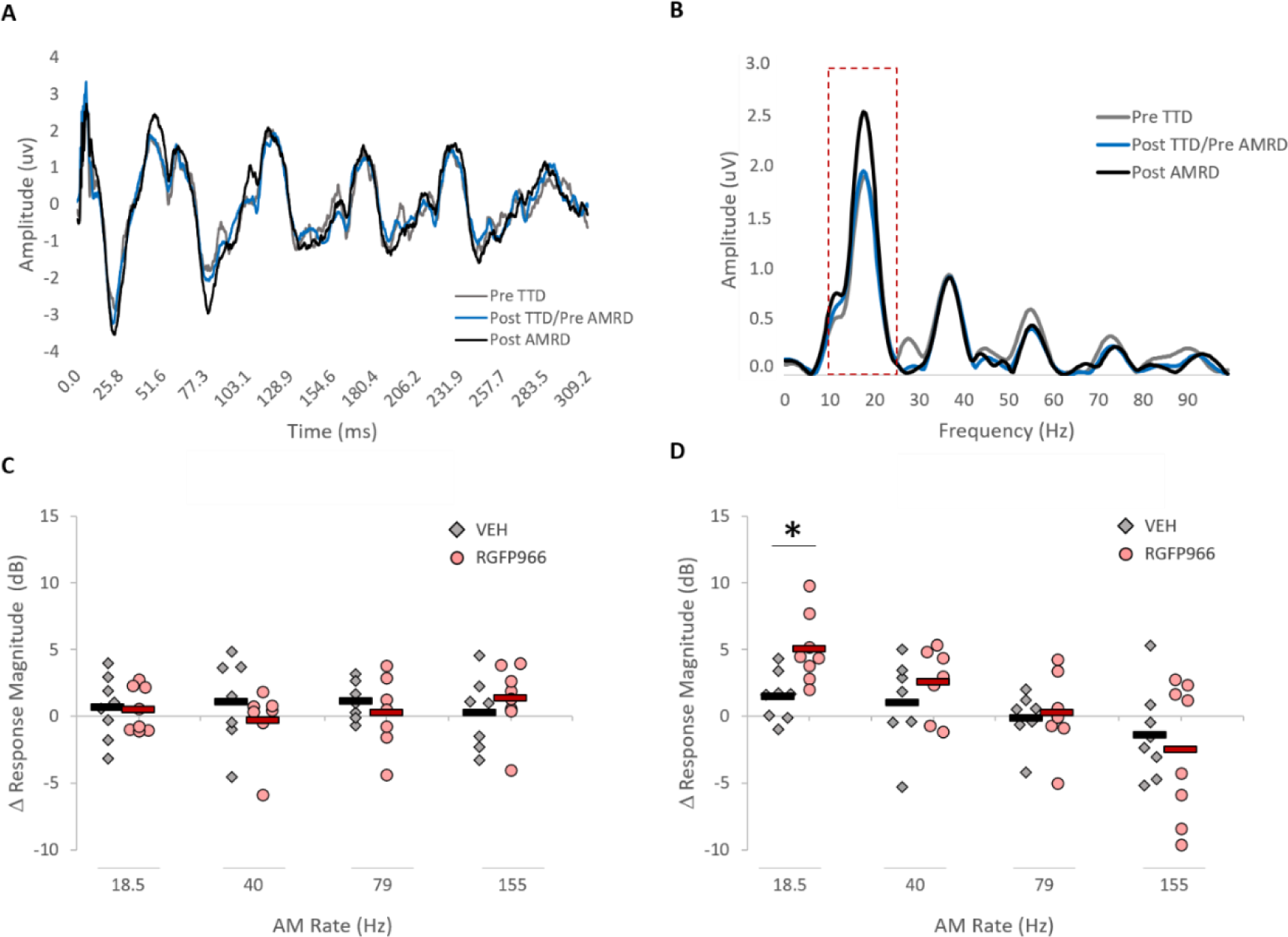
HDAC3 inhibition results in enhanced FFR response magnitude that is selective to the rewarded AM rate. (A) A representative FFR trace evoked by 18.5 Hz AM noise at the pre-tone discrimination training , post-tone discrimination training, and post-AM rate discrimination training time points in an RGFP966-treated rat.*. (B)* A fast Fourier transform of the FFR shown in panel A reveals stronger encoding of the 18.5 Hz AM rate following AM rate discrimination training. (C) Over the course of tone-tone discrimination leaning, response magnitude is stable in AM noise-evoked FFRs. (D) Over the course of AM rate discrimination learning, RGFP966-treated rats show a selective increase in response magnitude of FFRs evoked by the rewarded 18.5 Hz S+ (p<0.01), and a greater increase *vs.* vehicle-treated rats. There were no other significant changes in response magnitude or drug treatment group differences. *p<0.05 *vs.* vehicle

### 3.4 HDAC3 inhibition results in stimulus-specific and general plasticity effects in the primary auditory cortex

Memory for associations between specific sound features like a particular acoustic frequency and its link to potential for rewards, including the highly specific auditory memory enabled by HDAC3 inhibition, has been associated with signal-specific plasticity at the level of the primary auditory cortex (A1) (Recanzone, Schreiner, & Merzenich, 1993; Polley et al., 2006; Keeling et al., 2008; Bieszczad et al., 2010; Bieszczad & Weinberger, 2012; Bieszczad et al., 2015; Shang et al., 2019; Rotondo & Bieszczad, 2020; Rotondo & Bieszczad, 2021). Cortical representations of temporal features of sound can likewise be transformed by experience (Bao et al., 2004). Here, we sought to characterize changes in auditory cortical encoding of AM sounds associated with the unusually specific memory enabled by HDAC3 inhibition.

Electrophysiological recordings in A1 were following the AM Memory Test to compare A1 plasticity in the treated groups (RGFP966 vs. vehicle) and the untreated, naïve control group on measures of phase-locking and response consistency. Though evoked responses to AM sounds do not significantly differ as a function of cortical frequency tuning (Bao et al., 2004), we note that the distribution of best frequency of cortical recording sites in vehicle- and RGFP966-treated rats did not differ from naïve rats (vehicle vs. naïve: *X*^2^ (2, *192)* = 3.79, *p* = 0.149; RGFP966 vs. naïve: *X*^2^ (2, 249) = 2.91, *p* = 0.232) (Table 6). Thus, it is unlikely that frequency tuning properties of recording sites explain group differences in evoked responses to AM sounds.

Previous studies have reported a learning-induced increase in auditory cortical response consistency evoked by sound that generalizes across different sound stimuli, though these studies did not include an explicit behavioral test of memory specificity induced by the completion of training, thus precluding the possibility of attributing the generalized effect on cortical response variability with a generalized behavioral effect across sound (Leon et al., 2008; Von Trapp et al., 2016). Here it was predicted that RGFP966-treated rats would have an increase in responses consistency above and beyond their vehicle-treated counterparts, and that this form of plasticity would be specific to the responses evoked by the training sounds.

To determine AM noise-evoked response consistency in A1, we calculated the sum of point-to-centroid distances using a *k-means* clustering approach. Here, *lower* values mean *greater* response consistency (i.e. lower response variability). There was a significant effect of training: vehicle- and RGFP966 treated rats exhibited a significant increase in response consistency for nearly all stimuli (Table 7; Fig. 7). In support of our prediction, RGFP966-treated rats had greater response consistency than vehicle-treated rats. However, this effect was observed for nearly all stimuli. Together, this suggests that changes in response consistency represent a generalized change in cortical sound processing that is induced by training and is facilitated by HDAC3-inhibition.

**Figure 7.**
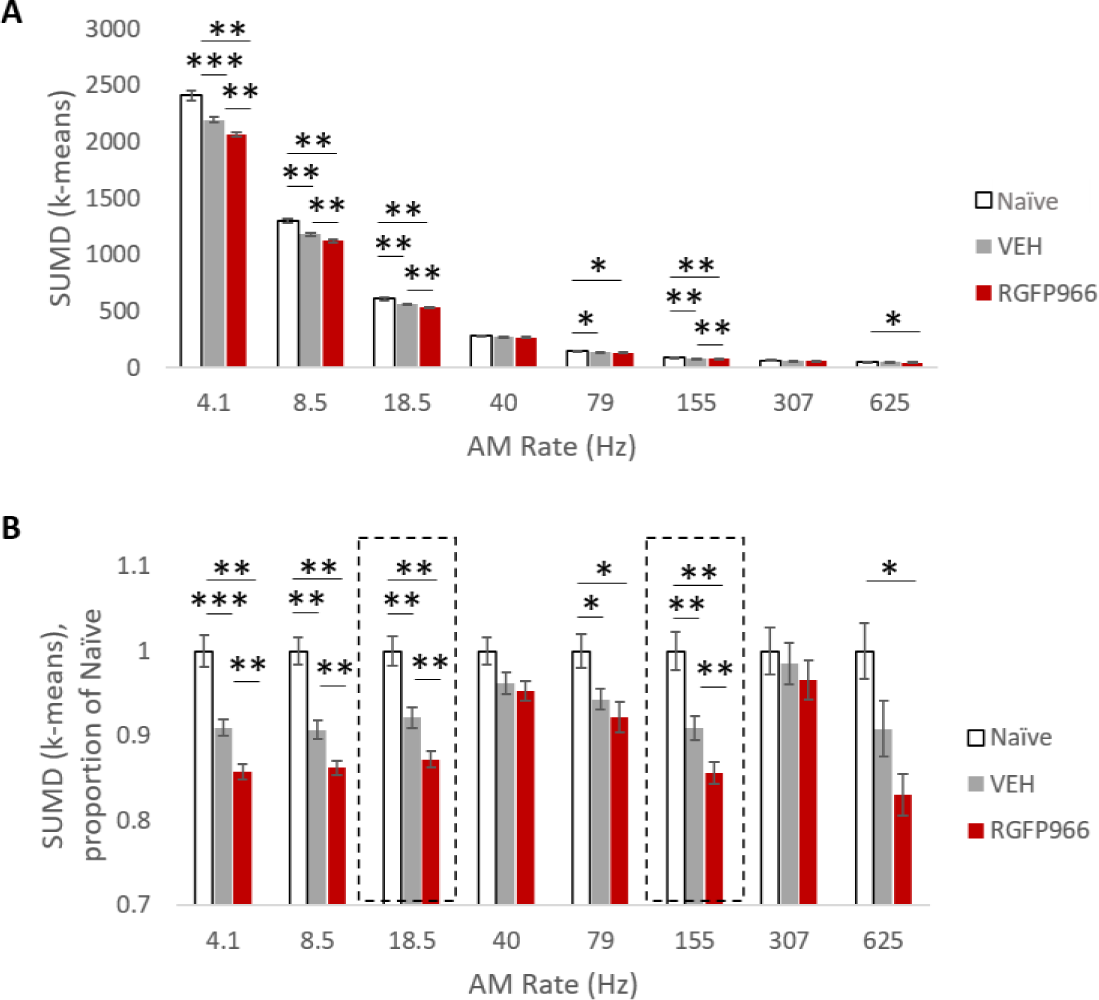
A1 response consistency is increased as a function of memory formation and of HDAC3i-enabled memory specificity. Panels display (A) the raw sum of point-centroid distances (SUMD) and (B) the sum of point-centroid distances (SUMD) as a proportion of naïve subjects. Response consistency increased with training, revealed by the lower SUMD in vehicle- and RGFP966-treated groups. Further, RGFP966-treated rats had greater response consistency than vehicle-treated rats. These effects generalized across most AM rates. *p<0.05, **p<0.01, ***p<0.001

**Table 6.** Distribution of best frequency (BF) for naïve, vehicle-treated, and RGFP966-treated rats. This table displays the percent of recording sites with a best frequency (BF) within approximately 2-octave bins for naïve, vehicle-treated, and RGFP966-treated rats.

| BF (kHz) | Naïve<br>(127 sites; 5 rats) | VEH<br>(192 sites; 8 rats) | RGFP966<br>(249 sites; 8 rats) |
| --- | --- | --- | --- |
| 0.59 - 2.37 | 34.67% | 36.45% | 38.15% |
| 3.36-13.45 | 41.73% | 45.83% | 42.57% |
| 19.02-53.81 | 23.62% | 17.71% | 19.27% |

**Table 7.**
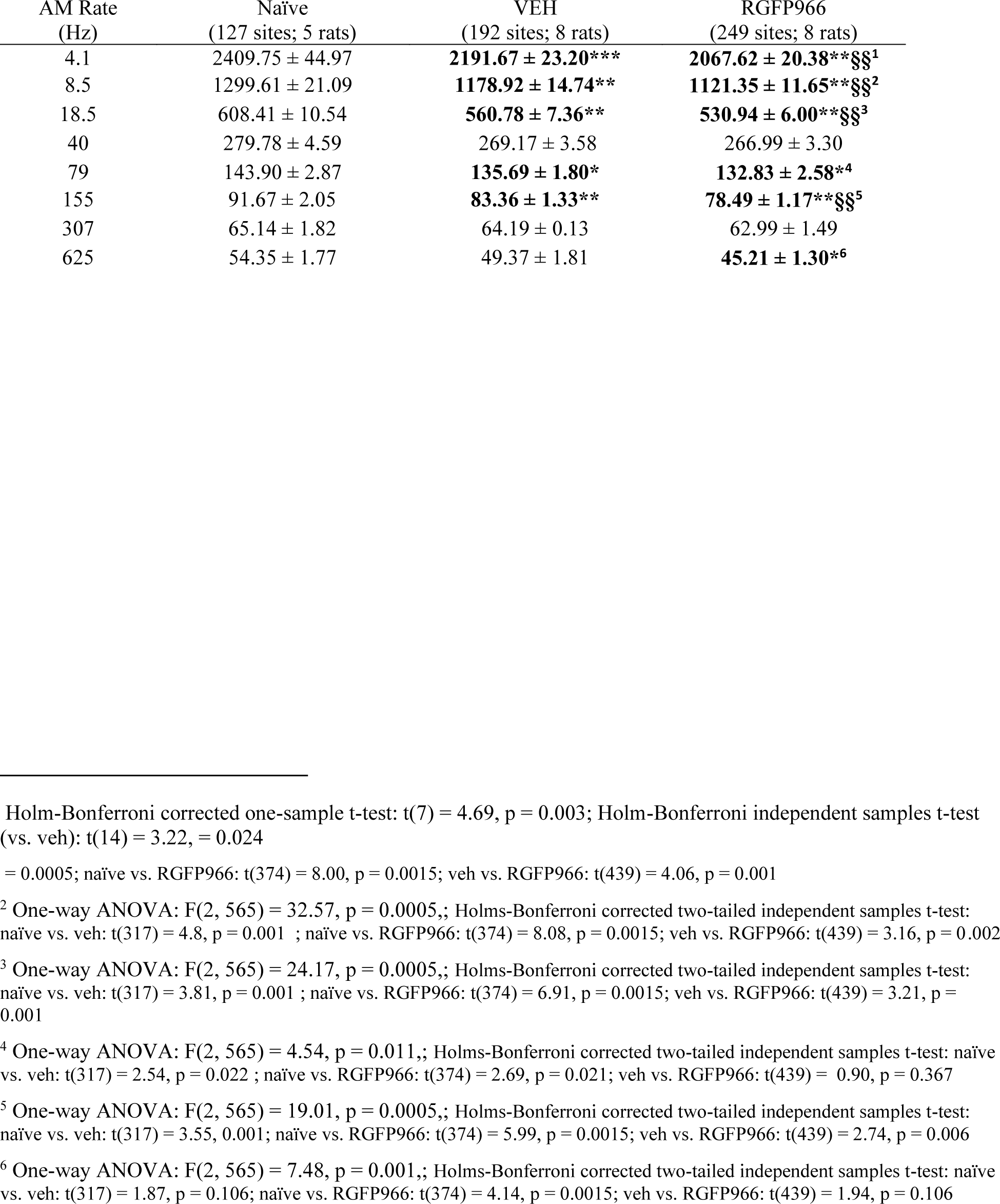
Within-stimulus response consistency in naïve, vehicle-treated, and RGFP966-treated animals. This table displays the sum of point-centroid distances determined by the *kmeans* clustering approach- a measure of within- stimulus response consistency. Not that *smaller* values indicate *greater* response consistency. Sample sizes are given in parentheses. All data are displayed as M ± SE. For VS data, significant differences are bolded. *indicates a difference vs. naïve animals, *p<0.05, **p<0.01, ***p<0.001. § indicates a difference vs. vehicle-treated animals, §§p<0.01

Previous studies have reported learning-induced increases the magnitude of phase-locked cortical activity, with some indirect evidence that changes in phase-locking may be stimulus specific (Beitel et al., 2003; Bao et al., 2004). To determine the strength of phase-locking, we calculated for each recording site within A1, a vector strength (VS) and the Rayleigh statistic (RS), which estimates the significance of VS controlling for the total number of spikes. At the cortical level of A1, the ceiling rates for phase-locking are substantially lower relative to brain structures closer to the periphery, which can follower faster rates of modulated sound (Joris et al., 2004). In anesthetized preparations, auditory cortical phase-locking is typically limited to AM rates below 20-30 Hz (Eggermont, 1991; Anderson et al., 2006; Fitzpatrick et al., 2009; Miller et al, 2002; Bao et al. 2004). Therefore, these analyses were restricted to a set of slower AM test rates: 4.1, 8.5, and 18.5 Hz (the S+).

Results for the phase-locking analyses differed from the response consistency analysis in two major ways. First, RGFP966-treated animals, but *not* vehicle-treated, animals showed learning-induced changes in phase-locking (Fig. 8; Table 8). Because both vehicle- and RGFP966-treated animals were able to learn the task equally, the differences in learning-induced phase-locking suggests that plasticity in cortical phase-locking may underlie a behavioral function beyond task acquisition that relates to the quality of the memory formed by learning (e.g., such as its specificity or strength). Second, changes in phase-locking exhibited a greater degree of stimulus specificity than response consistency. RGFP966-treated rats, compared to naïve or vehicle-treated rats, showed a significant increase in vector strength in responses evoked by the rewarded (S+) 18.5 Hz AM rate, but no changes in responses evoked by the neighboring, novel 8.5 Hz AM rate. Further, RGFP966-treated rats showed a significant *decrease* in vector strength in responses evoked by the novel 4.1 Hz AM rate. Together, this pattern of changes suggests a selective enhancement in phase-locked responses to the behaviorally relevant 18.5 Hz rate.

**Figure 8.**
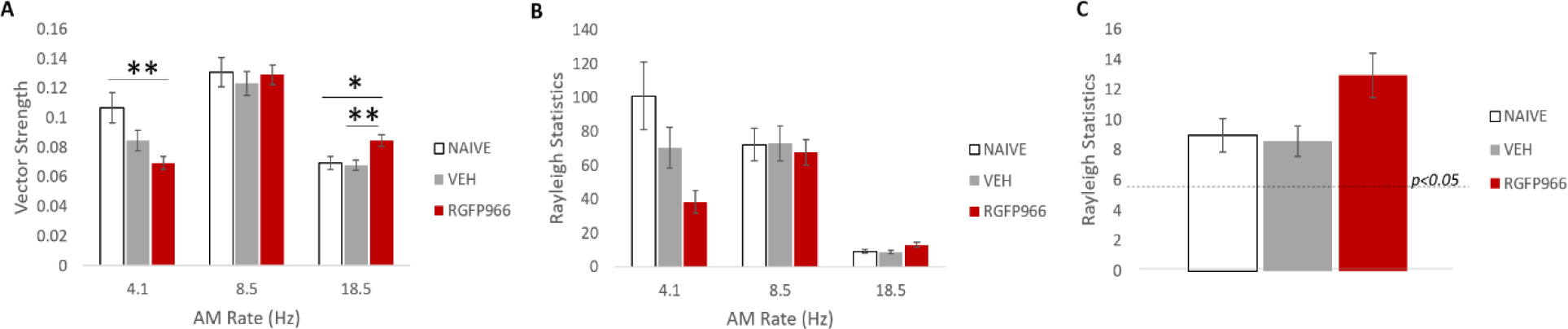
HDAC3 inhibition enhances A1 phase-locking to the rewarded AM rate. **(A)** Vector strength, a measure of phase-locking strength) is significantly greater in responses evoked by the rewarded 18.5 Hz AM rate among RGFP966 treated animals, vs. naïve and vehicle-treated animals. Vector strength is significantly lower in responses evoked by the novel 4.1 Hz AM rate among RGFP966 animals vs. naïve animals. **(B)** The Rayleigh statistic estimates the significance of phase-locking, where a value of 5.991 corresponds to p<0.05 and a value of 13.816 corresponds to p<0.001. **(C)** A zoomed-in view of the Rayleigh statistics for responses evoked by the rewarded 18.5 Hz AM rate. The dashed line represents the threshold value for significant phase-locking.

**Table 8.**
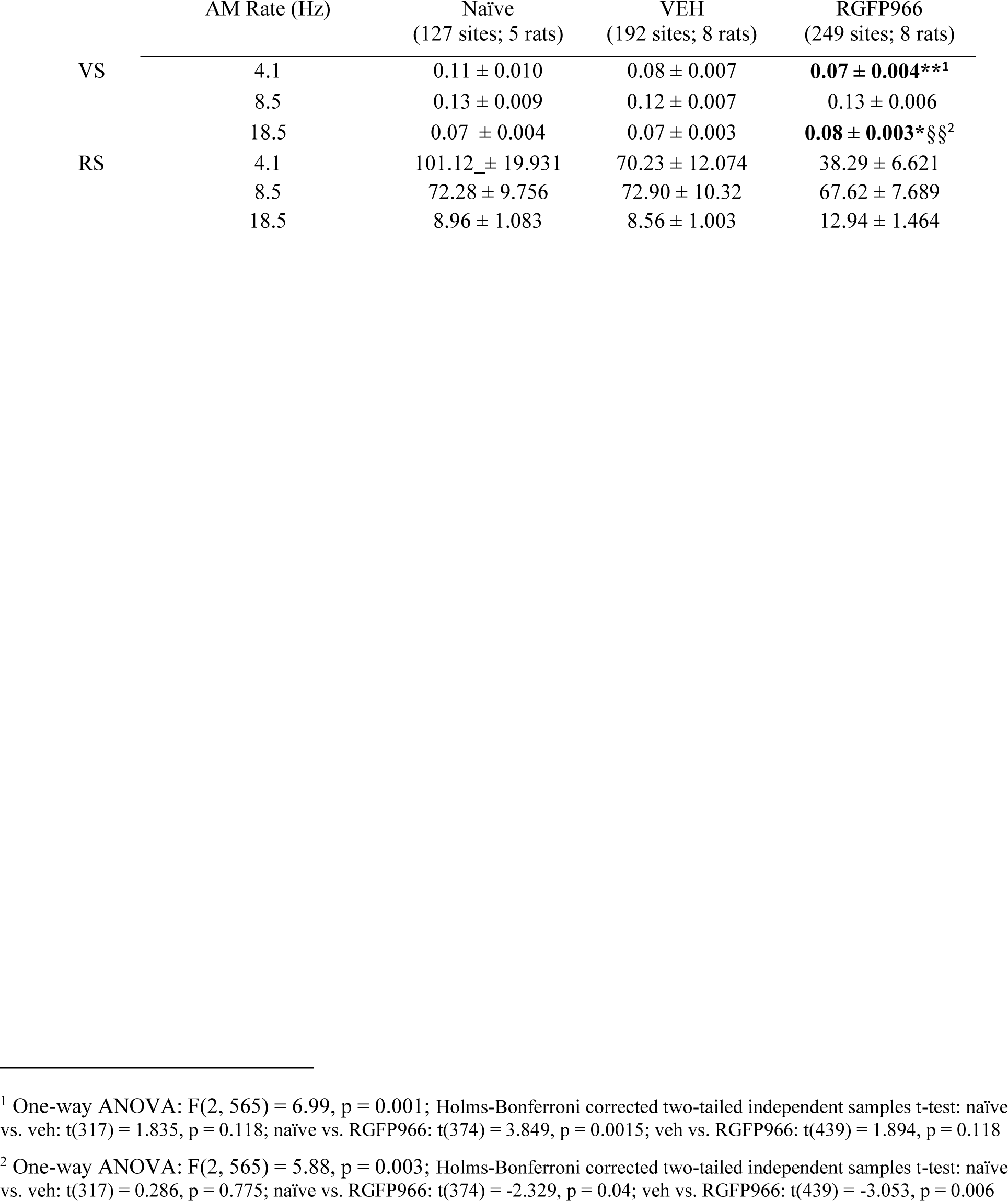
Vector strength and Rayleigh statistic in naïve, vehicle-treated, and HDAC3-treated animals. This table displays vector strength (VS) and the Rayleigh statistic (RS), which estimates the significance of VS taking into account the total number of spikes. Sample sizes are given in parentheses. All data are displayed as M ± SE. For VS data, significant differences are bolded. *indicates a difference vs. naïve animals, *p<0.05, **p<0.01. § indicates a difference vs. vehicle-treated animals, §§p<0.01 For RS data, the critical values are 5.991 for α = 0.05 and 13.816 for α = 0.001.

|  | AM Rate (Hz) | Naïve<br>(127 sites; 5 rats) | VEH<br>(192 sites; 8 rats) | RGFP966<br>(249 sites; 8 rats) |
| --- | --- | --- | --- | --- |
| VS | 4.1 | $0.11 \pm 0.010$ | $0.08 \pm 0.007$ | **$0.07 \pm 0.004^{**1}$** |
| | 8.5 | $0.13 \pm 0.009$ | $0.12 \pm 0.007$ | $0.13 \pm 0.006$ |
| | 18.5 | $0.07 \pm 0.004$ | $0.07 \pm 0.003$ | **$0.08 \pm 0.003^{*§§2}$** |
| RS | 4.1 | $101.12 \pm 19.931$ | $70.23 \pm 12.074$ | $38.29 \pm 6.621$ |
| | 8.5 | $72.28 \pm 9.756$ | $72.90 \pm 10.32$ | $67.62 \pm 7.689$ |
| | 18.5 | $8.96 \pm 1.083$ | $8.56 \pm 1.003$ | $12.94 \pm 1.464$ |

<sup>1</sup> One-way ANOVA: $F(2, 565) = 6.99$ , $p = 0.001$ ; Holms-Bonferroni corrected two-tailed independent samples t-test: naïve vs. veh: $t(317) = 1.835$ , $p = 0.118$ ; naïve vs. RGFP966: $t(374) = 3.849$ , $p = 0.0015$ ; veh vs. RGFP966: $t(439) = 1.894$ , $p = 0.118$
<sup>2</sup> One-way ANOVA: $F(2, 565) = 5.88$ , $p = 0.003$ ; Holms-Bonferroni corrected two-tailed independent samples t-test: naïve vs. veh: $t(317) = 0.286$ , $p = 0.775$ ; naïve vs. RGFP966: $t(374) = -2.329$ , $p = 0.04$ ; veh vs. RGFP966: $t(439) = -3.053$ , $p = 0.006$

To further investigate these changes in cortical phase-locking, we next determined the proportion of cortical recordings sites that exhibited significant phase-locked responses for each AM rate, using the criteria of a Rayleigh statistic ≥ 5.991 (Table 9; Fig. 9). RGFP966-treated rats exhibited an increased proportion of sites with significant phase-locking to the 18.5 Hz AM rate (0.461/ 115 of 249 cites), vs. naïve rats (0.377; 48 of 127 sites). In contrast, both RGFP966- (0.634; 158 of 249 sites) and vehicle-treated (0.625; 120 of 192 sites) rats exhibited a decreased proportion of sites with significant phase-locking to the 4.1 Hz AM rate, vs. naïve rats (0.795; 101 of 127 sites). Given that these proportional changes mimicked the pattern of changes in phase-locking (Fig. 8; Table 8), we next tested whether group differences in phase-locking would still be observed after removing the sites with no significant phase-locked response. When only considering sites with significant phase-locked responses, RGFP966-treated rats still had greater vector strength in responses evoked by the rewarded 18.5 Hz AM rate (Fig. 3.9c; Table 9) and weaker vector strength in responses evoked by the novel 4.1 Hz AM rate (Fig 3.9a; Table 9). Therefore, RGFP966 treated rats exhibit at least two distinct forms of plasticity related to phase-locking: (a) a change in the proportion of phase-locked sites and (b) a change in the strength of phase-locking both overall, and within just the phase-locked sites. This pattern of changes may also reflect “competitive loss,” (here, an increase in metabolic resources dedicated to representing behavioral relevant sounds at the expense of novel, distinct sounds) especially when behaviorally relevant sounds with enhanced phase-locking are near the ceiling of cortical phase-locking ability (i.e. 20-30 Hz) (Bao et al., 2004).

**Figure 9.**
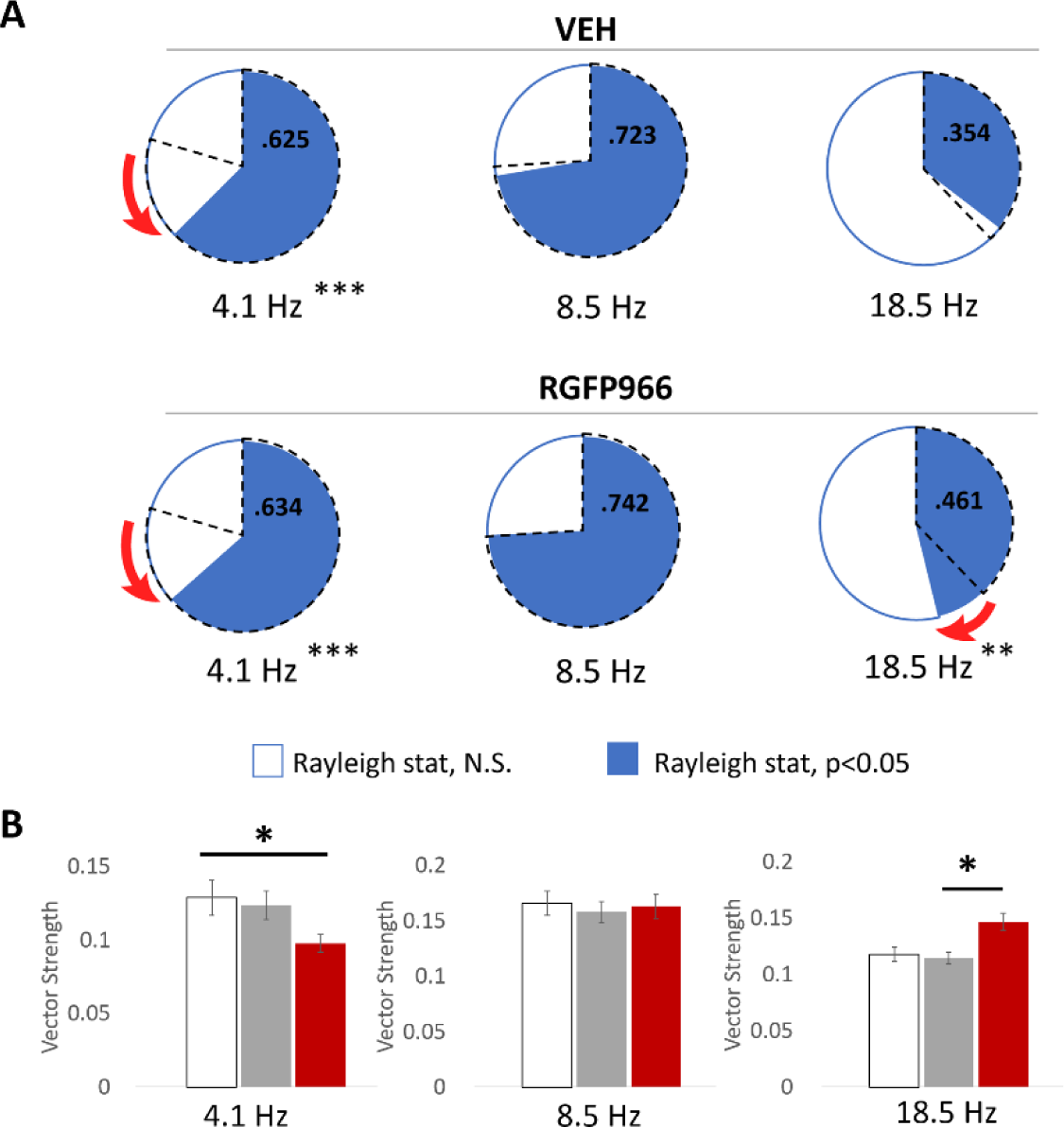
HDAC3 inhibition increases the proportion of cortical sites that phase-locked to the rewarded 18.5 Hz AM rate and the strength of phase-locking within those sites. **(A)**The pie charts displays the proportion of recordings sites in A1 in vehicle- and RGFP966-treated animals with a Rayleigh statistic ≥ 5.991, which corresponds to p < 0.05. The dashed lines represent the proportion of cortical sites with significant phase-locked responses in naïve animals. Red arrows represent the direction of significant changes. **p<0.01 vs. naïve; ***p<0.001 vs. naïve **(B)** When only considering cortical sites with a Rayleigh statistic ≥ 5.991, RGFP966-treated animals have significantly greater vector strength in responses evoked by the rewarded 18.5 Hz and significantly weaker vector strength in responses evoked by the novel 4.1 Hz, vs. naïve rats. *p<0.05

**Table 9.**
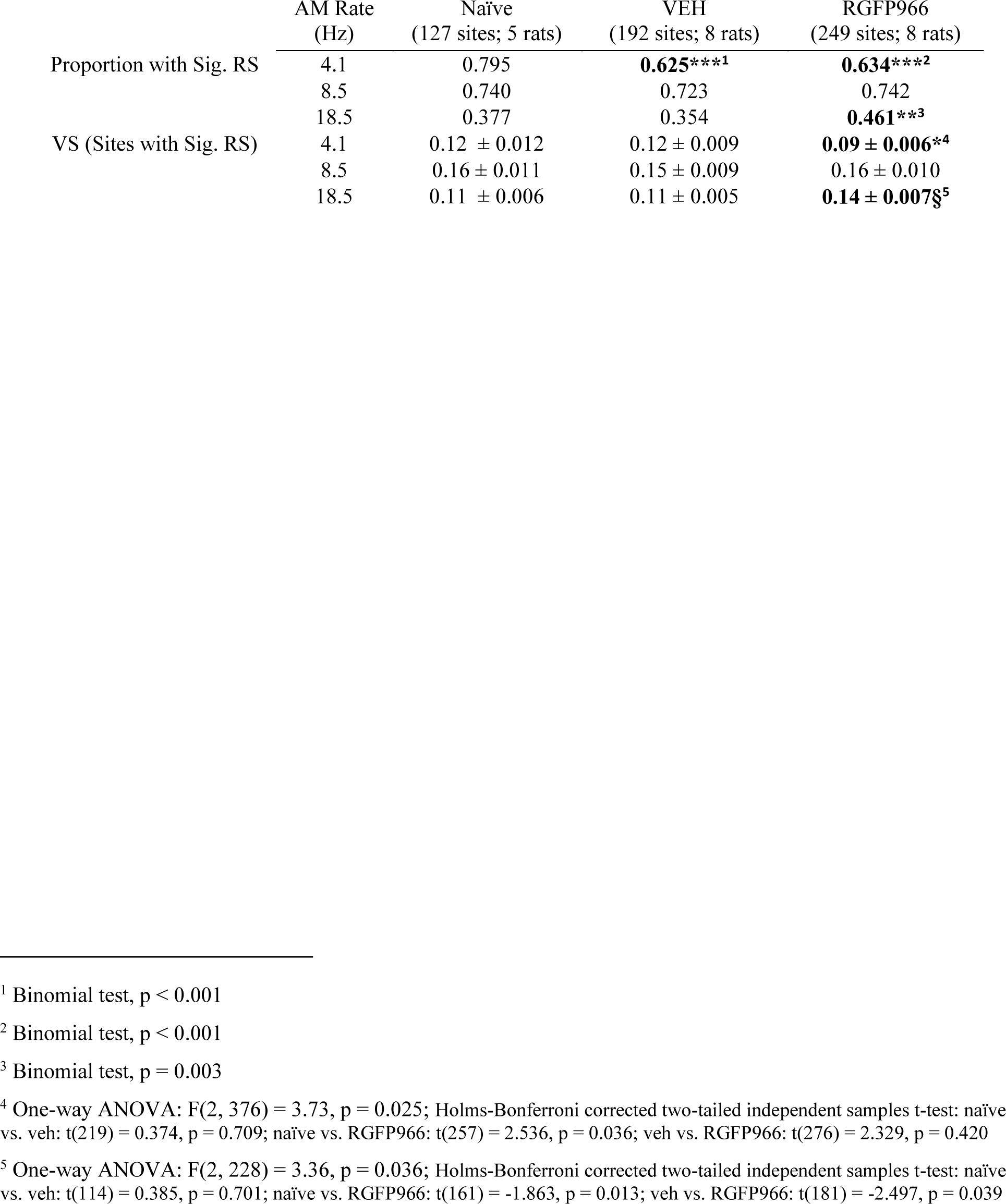
Vector strength in cortical sites with significant phase-locking in naïve, vehicle-treated, and HDAC3-treated animals. This table shows the proportion of cortical sites with significant phase-locking (p<0.05), as revealed by a Rayleigh statistics (RS) value of at least 5.991. Vector strength (VS) for sites with significant phase-locking is displayed (M ± SE). Significant differences are bolded. *indicates a difference vs. naïve animals, *p<0.05, **p<0.01, ***p<0.001. § indicates a difference vs. vehicle-treated animals, §p<0.05

**Table 10.** Summary of correlations with phase-locked neural responses and sound-cued behavior. This table displays the Pearson r Values for correlations between phase-locked neural responses measured from the scalp with the FFR (response magnitude [Resp. Mag.]) or recorded extracellularly from A1 (vector strength [VS], Rayleigh statistic [RS], % of phase-locked sites) and the percent of responses to the rewarded 18.5 Hz AM rate (% BPs to S+) during the AM rate memory test). Δ Resp. Mag refers to the difference in FFR response magnitude pre-to-post AM rate discrimination training. Sample sizes are in parentheses and remain consistent across analyses except for vector strength among phase-locked units. Because one rat had 0 phase-locked sites, there was no data point to include in this analysis. Significant correlations are bolded. *p<0.05, **p<0.01, ***p<0.001

|  | All | VEH | RGFP966 |
| --- | --- | --- | --- |
| VS vs. $\Delta$ Resp. Mag. | **0.837\*\*\***<br>(16) | 0.685<br>(8) | **0.819\***<br>(8) |
| RS vs. $\Delta$ Resp. Mag. | **0.891\*\*\***<br>(16) | **0.780\***<br>(8) | **0.911\*\***<br>(8) |
| VS vs. % BPs to S+ | **0.684\*\***<br>(16) | 0.393<br>(8) | 0.648<br>(8) |
| RS vs. % BPs to S+ | **0.768\*\*\***<br>(16) | 0.313<br>(8) | **0.804\***<br>(8) |
| % Phase-Locked Sites vs. %BPs | **0.525\***<br>(16) | 0.268<br>(8) | 0.443<br>(8) |
| VS (for Phase-Locked Sites) vs. % BPs | **0.721\*\***<br>(15) | 0.402<br>(7) | **0.731\***<br>(8) |
| $\Delta$ Resp. Mag vs. % BPs | **0.755\*\*\***<br>(16) | 0.156<br>(8) | **0.740\***<br>(8) |

Overall, there is a complex pattern of cortical changes that occur with memory for AM sounds and may be modified by the level of specificity with which memory is formed. Response consistency for most AM rates is improved with training, with more pronounced effects in the HDAC3-inhibited group vs. vehicle-treated group. Interestingly, there is a significant relationship between response consistency and the Rayleigh statistic (18.5 Hz: n = 568, r = -0.226, p <0.00001; 8.5 Hz: n = 568, r = -0.272, p = <0.00001; 4.1 Hz: n = 568, r = -0.074, p = 0.078), suggesting that better response consistency may *facilitate* significant phase locking, as has been previously proposed (White-Schwoch et al., 2017). However, because better response consistency was also found in instances with weaker phase locking at the group level (as in responses evoked by 4.1 Hz in the vehicle- and RGFP966-tretaed groups), this does not appear to be a causal relationship between response consistency and the strength of phase locking. Similarly, changes in the proportion of cortical sites that phase-lock to AM rates and changes to the strength of phase-locking appear to be partially dissociated at the level of the group, suggesting that recruitment of neurons into phase-locked responses and the strength of phase-locked responses are two distinct phenomena. Nonetheless, all observed forms of plasticity are more likely to occur with HDAC3 treatment and the formation of highly specific memory. However, it is important to note that plasticity related to phase-locking measures, but *not* response consistency measures, was signal-specific. Therefore, we followed up by testing whether these signal-specific forms of phase-locking plasticity (as oppose to the other form discovered) were the substrates of behavioral memory specificity.

### 3.5 Auditory system plasticity is correlated with individual differences in memory specificity

To validate forms of signal-specific auditory plasticity as substrates of memory specificity for AM rate, we pursued correlations between neural measures sensitive to phase-locking and behavioral measures that index memory specificity. While there were multiple potential indices of memory specificity, we focused on the percent of responses to the 18.5 Hz S+ during the memory test for several reasons. First, memory specificity effects were largely driven by differences in responding to the S+, as opposed to the S- (Fig. 5; Table 4). Further, the percent of responses to the S+ is a straightforward metric that is highly correlated with other specificity indices (% BPs to S+ vs. Δ % BPs to S+ vs. S-: r = 0.981, p <0.00001; vs. Δ % BPs to S+ vs. nearby neighbors: r = 0.987, p <0.00001; vs. Δ % BPs to S+ vs. distant neighbors: r = 0.976, p < 0.00001).

It was first important to understand whether the magnitude of phase-locked neural activity measured in the FFR was correlated with phase-locked activity measured in the auditory cortex. As noted previously (see: *2.4 Auditory Brainstem Response Recordings and Analysis),* FFRs evoked by the 18.5 Hz S+ likely include a significant cortical component, in addition to subcortical components. Indeed, the learning-induced change in FFR response magnitude was significantly correlated with phase-locking strength in the auditory cortical recordings (Fig. 10a; Table 10). While the data does not exclude the possibility of subcortical plasticity, we cannot tease apart cortically- vs subcortically-sourced components of the FFR data. Notably, stronger phase-locking in responses evoked by 18.5 Hz, whether measured from the FFRs (Fig. 10b; Table 10) or from cortical recordings (Fig. 10c,d; Table 10) was associated with a greater degree of memory specificity.

**Fig. 3.10.**
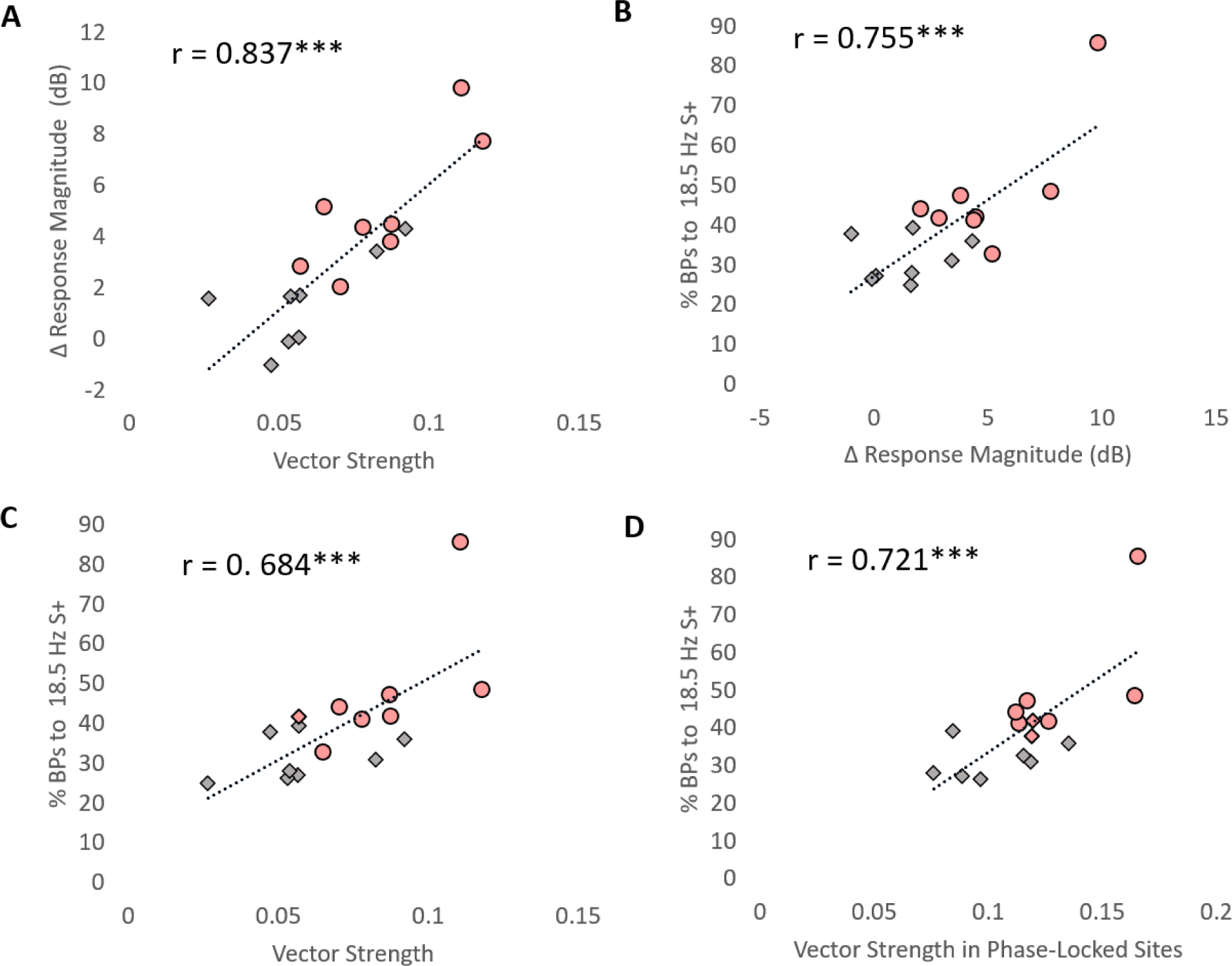
Auditory system plasticity is correlated with AM-rate specific memory. Each panel presents correlative data between two of several different types of measures: auditory cortical phase locking to the 18.5 Hz AM rate (vector strength), learning-induced change in FFR response magnitude evoked by the 18.5 Hz AM rate, and the percent of behavioral responses to the 18.5 Hz S+ during the AM rate memory test. **(A)** Greater A1 phase-locking to 18.5 Hz is related to greater increases in FFR response magnitude to 18.5 Hz. **(B)** A greater increase in response magnitude in FFRs by the 18.5 Hz S+ is related to a to a greater degree of memory specificity, as indexed by the percent of behavioral responses to 18.5 Hz S+ during the AM rate memory test. Stronger auditory cortical phase- locking, when considering all recording sites in A1 **(C)** or *only* responses from phase-locked sites **(D)** is also related to a greater degree of memory specificity.

## 4. DISCUSSION

We report for the first time support that HDAC3 inhibition (HDAC3i) enhances behavioral memory specificity for a temporal feature of sound, amplitude modulation (AM) rate. Greater memory specificity enabled by HDAC3i was characterized by several learning-induced forms of neurophysiological plasticity in the primary auditory cortex (A1). Unlike vehicle-treated rats, HDAC3i rats exhibited an increase in the magnitude of phase-locking that was *specific* to the rewarded AM rate- an effect that was also validated by AM rate-specific effects in the surface recorded frequency following response (FFR). HDAC3i rats also showed a larger increase in A1 response consistency (relative to vehicle-treated rates) that generalized across responses evoked by a range of AM stimuli. Further analysis revealed that the cortical increase in sound-specific phase-locking was attributable to both an increase in the proportion of recording sites that participate in the phase-locked response and an increase in the strength of phase locking among these sites (i.e. after removing sites that do not exhibit significant phase-locking). Brain-behavior relationships revealed that measures of phase-locking correlated with individual differences in behavioral memory specificity among HDAC3i- and vehicle-treated rats. In sum, these findings support the that the effect of HDAC3i to enhance memory specificity may be mediated by enabling signal-specific auditory neuroplasticity. The findings extend this hypothesis for the first time to memory formation for *temporal* features of sounds with *temporal coding* strategies in the auditory brain.

### 4.1. HDAC3 inhibition enhances temporal coding of the rewarded sound

Temporal coding, via phase-locking to sound features, can increase the precision of sound-evoked neural activity behaviorally relevant information (Lakatos et al., 2008; Schroeder et al., 2010; Peelle, Gross, & Davis, 2013). Here, we report an increase in the strength of phase-locking to the rewarded AM rate in the primary auditory cortex among HDAC3i rats with unusually specific memory. Notably, when considering only recording sites with significant phase-locked responses, the pattern of effects with respect to strength of phase-locking hold: HDAC3i rats exhibit enhanced phase-locking to the rewarded AM rate. This suggests that the highly specific memory for AM rate entails both a change in the neurons that are recruited into the response *and* a change in the response characteristics within neurons (Bao et al., 2004). This signal-specific enhancement of cortical phase-locking is in line with findings that phase-locking is shaped by both acoustic information *and* other non-sensory and experience-dependent information such as meaning (Bao et al., 2004; Strait et al., 2012; Peelle et al., 2013). Consistent with this, we report that individual differences in the strength of phase-locking were significantly correlated with the degree of memory specificity for the rewarded sound.

The observation of enhanced cortical phase-locking to the rewarded AM rate in extracellular cortical recordings was validated by the FFR, which is a signal with both cortical and subcortical components (Coffey et al., 2016). Indeed, measures of the magnitude of phase-locked activity in the FFR with magnitude of phase-locked activity in the primary auditory cortex were significantly positively correlated. Nonetheless, the FFR was recorded chronically, and revealed that phase-locking is enhanced within individual subjects over the course of learning about AM sounds, and even more so in animals with highly specific memory for the rewarded AM sound.

Interestingly, the FFR did not reveal a significant change in phase-locking evoked by the explicitly unrewarded AM sound (155 Hz), even though this rate is within the range of subcortical phase locking ability (Joris et al., 2004; Fitzpatrick et al., 2009; Coffey et al., 2016). It is possible that this form of plasticity is not readily induced at the subcortical level. Alternatively, because 18.5 Hz and 155 Hz are on different sides of a perceptual barrier, such that modulations at 18.5 Hz are perceived as “flutter” while modulations at 155 Hz are perceived as “roughness” or even tonal (Besser, 1967; Krumbholz, Patterson, & Pressnitzer, 2000), animals could use the sound quality, rather than specific differences in AM rate, to guide their behavioral responses. Indeed, this may be consistent with the pattern of results observed in the vehicle-treated group, which displayed an observable degree of memory specificity despite not showing any significant enhancements in phase-locking. If this is true, then it is notable that treatment with HDAC3i may bias subjects to encode specific features of the stimulus even when they are not strictly necessary to solve the task (i.e. since other strategies could have been used), as has been suggested by previous results (Bieszczad et al., 2015; Rotondo & Bieszczad, 2020; Rotondo & Bieszczad, 2021).

### 4.2 HDAC3 inhibition enhances the learning-induced increase in auditory cortical response consistency to AM sounds

In contrast to the signal-specific changes in phase-locking, we observed another form of plasticity with a temporal component that instead generalized across AM sounds. Specifically, we observed an increase in sound-evoked response consistency (i.e. a decrease in response variability) in the primary auditory cortex among both trained groups, with a significantly greater increase in HDAC3i rats. That auditory training improves auditory response consistency, even for sounds that were not explicitly trained, is in line with previous studies (Leon et al., 2008; Hornickel et al., 2012; Anderson et al., 2013).

Many studies have correlated auditory-evoked response consistency with proficiency in auditory and language skills (Anderson et al., 2012; Hornickel & Kraus, 2013; Von Trapp et al., 2016; White-Schwoch et al., 2017; Caras & Sanes, 2019). Thus, one interpretation of this data is that response consistency may *facilitate* learning within the AM category, as it could provide a foundation of consistently discriminable neural representations of acoustically similar sounds that facilitates behavioral discrimination between them. Indeed, subjects with the best response consistency also had enhancements in phase-locking and a greater degree of memory specificity, though these measures were not consistently correlated. Future analyses will consider the potential interactive contribution of these forms of plasticity to learned behavior.

As with phase-locking, response consistency may be improved with attention or task-engagement (Von Trapp et al., 2016). However, these factors are not a prerequisite, as other studies, including the present one, have observed improved response consistency in absence of attentional or contextual factors (Leon et al., 2008) or active task engagement (Hornickel et al., 2012; Anderson et al., 2013). Given that HDAC3i enhanced response consistency above and beyond the level of vehicle-treated controls, and that these effects were observed approximately 3 days after the final auditory training session, these are long-lasting effects that may be due to a change in gene expression regulated by HDAC3. Indeed, certain genes have been implicated in auditory cortical (Centanni et al., 2014) and subcortical (Selinger et al., 2016) temporal response consistency. HDAC3i may facilitate or prolong expression of genes critical to temporal encoding of sounds that result in enhanced physiological plasticity and a greater degree of behavioral memory specificity.

In sum, epigenetic manipulations, like HDAC3i, support the formation of unusually specific memory for temporal features of sound. Moreover, for the first time, we demonstrate that HDAC3i can alter temporal coding of sound features in the auditory brain. Like memory specific to acoustic frequency, memory specific to AM rate is supported by a signal-specific pattern of neurophysiological changes that provide discriminable representations of behaviorally relevant sound features. These neurophysiological changes appear to be susceptible to HDAC3-regulation, regardless of the stimuli evoking those patterns of responses, or the neural coding strategy (Bieszczad et al., 2015; Rotondo & Bieszczad, 2020; Rotondo & Bieszczad, 2021). Together, this supports that epigenetic regulators, like HDAC3, play an important role in the development of precise representations of spectral and temporal features of sound, both of which are critical for communication skills. More broadly, it supports a hypothesis in which HDAC3i can target and transform the detailed, “in the moment” sensory information received by the brain into long-term storage in memory, regardless of the type of experience or the way it is encoded in the brain (Bieszczad et al., 2015; Phan & Bieszczad, 2016).

## ACKNOWLEDGEMENTS

This work was supported by NIH R03-DC014753 (to K.M.B.) NIH R01-DC018561 (to K.M.B.), the American Speech-Hearing-Language Foundation 2017 New Century Scholars Grant (to K.M.B.) and The Brain & Behavior Foundation 2017 NARSAD Young Investigator Award (to K.M.B.).

